# Coordination of human hippocampal sharpwave-ripples during NREM sleep with cortical theta bursts, spindles, downstates and upstates

**DOI:** 10.1101/702928

**Authors:** Xi Jiang, Jorge Gonzalez-Martinez, Eric Halgren

**Author notes:** Submitting Author (Xi Jiang). Corresponding Authors (1-845-857-6020), (1-858-822-8385).

## Abstract

In rodents, waking firing patterns replay in NREM sleep during hippocampal sharpwave-ripples (HC-SWR), correlated with neocortical graphoelements (NC-GE). NC-GE include theta-bursts, spindles, downstates and upstates. In humans, consolidation during sleep is correlated with scalp-recorded spindles and down-upstates, but HC-SWR cannot be recorded non-invasively. Here we show in humans of both sexes that HC-SWR are highly correlated with NC-GE during NREM, with significantly more related HC-SWR/NC-GE for downstates or upstates than theta-bursts or spindles, in N2 than N3, in posterior than anterior HC, in frontal than occipital cortex, and ipsilaterally than contralaterally. The preferences interacted, e.g. frontal spindles co-occurred frequently with posterior HC-SWR in N2. These preferred GE, stages and locations for HC-SWR/NC-GE interactions may index selective consolidation activity, although that was not tested in this study. SWR recorded in different HC regions seldom co-occurred, and were related to GE in different cortical areas, showing that HC-NC interact in multiple transient, widespread but discrete, networks. NC-GE tend to occur with consistent temporal relationships to HC-SWR, and to each other. Cortical theta-bursts usually precede HC-SWR, where they may help define cortical input triggering HC-SWR firing. HC-SWR often follow cortical downstate onsets, surrounded by locally-decreased broadband power, suggesting a mechanism synchronizing cortical, thalamic and hippocampal activities. Widespread cortical upstates and spindles follow HC-SWR, consistent with the hypothesized contribution by hippocampal firing during HC-SWR to cortical firing-patterns during upstates and spindles. Overall, our results describe how hippocampal and cortical oscillations are coordinated in humans during events that are critical for memory consolidation in rodents.

**Significance Statement:** Hippocampal sharpwave-ripples, essential for memory consolidation, mark when hippocampal neurons replay waking firing patterns. In rodents, cortical sleep waves coordinate the transfer of temporary hippocampal to permanent cortical memories, but their relationship with human HC-SWR remains unclear. We show that human hippocampal sharpwave-ripples co-occur with all varieties of cortical sleep waves, in all cortical regions, and in all stages of Non-REM sleep but with overall preferences for each of these. We found that sharpwave-ripples in different parts of the hippocampus usually occurred independently of each other, and preferentially interacted with different cortical areas. We found that sharpwave-ripples typically occur after certain types of cortical waves, and before others, suggesting how the cortico-hippocampo-cortical interaction may be organized in time and space.

## Introduction

Neocortical declarative memory consolidation depends on hippocampal input (Squire et al., 2001) during non-rapid eye-movement (NREM) sleep (Rasch and Born, 2013). NREM sleep is characterized by large stereotyped neocortical (NC) population events (“graphoelements”, or GE) including theta-bursts (TB) (Gonzalez et al., 2018), sleep-spindles (SS) (Mak-McCully et al., 2017), downstates (DS) (Cash et al., 2009), and upstates (US) (Sanchez-Vives and McCormick, 2000). The hippocampus (HC) generates sharpwave-ripples (SWR) during NREM: a large negative ∼50-100 ms “sharp wave” with superimposed 80-200 Hz “ripples”, followed by ∼200 ms positive wave (Buzsáki, 2015). During SWR, rodent HC pyramidal cells “replay” spatio-temporal firing patterns from waking (Wilson and McNaughton, 1994), and coordinated replay occurs in visual (Ji and Wilson, 2007) and prefrontal (Peyrache et al., 2009; Johnson et al., 2010) cortices. Thus, hippocampal replay during SWR could inform neocortical firing patterns which directly underlie consolidation in animals (Buzsáki, 2015). Furthermore, the co-occurrence of NC-GE with HC-SWR in NREM sleep may help coordinate consolidation (Siapas and Wilson, 1998; Mölle et al., 2006). Indeed, disrupting hippocampal replay (de Lavilléon et al., 2015), HC-SWR (Ego-Stengel and Wilson, 2010; Suh et al., 2013; Rothschild et al., 2016), or NC-SS—especially when they occur with DS-US (Schreiner and Rasch, 2015; Maingret et al., 2016; Latchoumane et al., 2017)—impair consolidation.

While the experiments above were in rodents, NC-GE and HC-SWR occur in all examined mammals (Buzsáki et al., 2013). In humans, SS, DS-US, and especially their co-occurrences recorded with scalp-EEG correlate with successful memory consolidation (Niknazar et al., 2015), suggesting that if HC-SWR correlated with NC-GE in humans, they may also support consolidation. However, although HC-SWR (Bragin et al., 1999; Staba et al., 2004; Axmacher et al., 2008; Le Van Quyen et al., 2008) and NC-GE (Csercsa et al., 2010; Nir et al., 2011; Piantoni et al., 2017) have been recorded with intracranial electrodes in humans, little is known about their relationship. Clemens *et al*. (2007, 2011) found that events possibly related to HC-SWR (mixed interictal spikes and ripples recorded near the parahippocampal gyrus) co-occurred with scalp-recorded SS. Similarly, Staresina *et al*. (2015) found a 5% increase in HC ripple-band power within 250 ms of scalp-recorded SS centers. Here we report the relationship between human NC-GE and HC-SWR, simultaneously recorded from within their respective structures.

Human HC-SWR were detected with strict requirements for the rodent-like waveform (i.e. multiple oscillations occurring in a systematic relationship to the sharpwave), distinguished from epileptiform activity, and then were related to NC-TB, SS, DS and US. Ventral versus dorsal HC in rodents and anterior versus posterior HC in humans have distinct anatomical connections and functional properties (Strange et al., 2014; Hrybouski et al., 2019). Furthermore, NREM sleep stages N2 and N3 are highly distinct, with SS and isolated DS-US sequences prominent in N2, and sustained DS-US oscillations characterizing N3 (Silber et al., 2007). Thus, we characterize the SWR-GE relationship separately for anterior vs posterior HC, different cortical areas, and different sleep stages.

We demonstrate widespread co-occurrence of NC-GE with HC-SWR for all GE types, NREM stages, and cortical and hippocampal regions, but with strong quantitative differences. The specific NC sites whose GE co-occurred with each HC site varied significantly between HC sites and GE types, indicating that, if HC-SWR↔NC-GE co-occurrences index consolidation (an unproved proposition in humans), then each involves a substantial but limited subset of the HC and NC. As in cortico-thalamic recordings, NC-GE surrounding HC-SWR tended to occur in the order TB→DS→SS/US. We found that NC-TB typically precede HC-SWR, and so may reflect the cortical input that influences the content of HC-SWR firing. HC-SWR were preceded by depressed broadband activity, and were increased following multiple NC-DS, suggesting that NC-DS may have a role in synchronizing NC and HC. Finally, HC-SWR are positioned to provide cortical input as it becomes activated during the NC-SS and NC-US. We thereby define the HC↔NC interaction in humans during events that are crucial for memory consolidation in rodents.

## Methods

### Overview

Hippocampal sharp-wave ripples (HC-SWR) were identified based on stringent criteria regarding their anatomical origin, frequency, and waveform, and separated from epileptiform activity using multiple criteria. The spectral characteristics and spatio-temporal distribution of HC-SWR were then characterized. Peri-stimulus time histograms were constructed of neocortical graphoelement (NC-GE) occurrence times with respect to the HC-SWR. Separate histograms were constructed for each: HC-NC channel pair, each GE (TB, SS, DS, and US), and for each sleep-stage (N2 or N3). Significant co-occurrences were tabulated, their peak latencies determined, and collated across HC-NC pairs. Co-occurrence frequency and anatomical pattern were examined across GE type, HC location (anterior versus posterior), sleep stage, and their combinations.

### Patient selection

Patients with long-standing drug-resistant partial seizures underwent SEEG depth electrode implantation in order to localize seizure onset and thus direct surgical treatment (see Table 1 for demographic and clinical information). Patients were from 16 to 60 years old, with globally typical SEEG rhythms in most channels (i.e., absence of diffuse slowing, widespread interictal discharges, highly frequent seizures, etc.) with no previous excision of brain tissue or other gross pathology, and with at least one HC contact in an HC not involved in the initiation of seizures. The resulting group of 20 patients includes 6 patients with an HC contact in a location with no interictal spikes, which were used to guide the protocols applied to the remaining 14 patients (Table 1). The 20 patients included 7 males, aged 29.8±11.9 years old (range 16-58). Electrode targets and implantation durations were chosen entirely on clinical grounds (Gonzalez-Martinez et al., 2013). All patients gave fully informed consent for data usage as monitored by the local Institutional Review Board, in accordance with clinical guidelines and regulations at Cleveland Clinic.

**Table 1.** Patient characteristics. Pt: patient. L: Left. R: Right. Dur.: duration. Std dev: standard deviation.

| Pt# | Age | Sex | Handedness | L aHC | L pHC | R aHC | R pHC | # Cortical Channels | HC Seizure-free? | Interictal-free? | # Sleep Periods | mean duration of | Std dev NREM (h) | Total N2 Dur. (h) | Total N3 Dur. (h) | SWR Rate, N2 aHC | SWR Rate, N2 pHC | SWR Rate, N3 aHC | SWR Rate, N3 pHC |
| --- | --- | --- | --- | --- | --- | --- | --- | --- | --- | --- | --- | --- | --- | --- | --- | --- | --- | --- | --- |
| 1 | 20 | M | R | x |  |  |  | 19 | Y | N | 3 | 2.2 | 0.78 | 2.8 | 3.3 | 11 |  | 19 |  |
| 2 | 51 | F | R | x |  |  |  | 12 | Y | N | 4 | 7.5 | 1.87 | 27.3 | 0 | 1.6 |  |  |  |
| 3 | 58 | F | R | x | x |  |  | 24 | Y | N | 4 | 7.1 | 2.44 | 23.1 | 0.30 | 11 | 12 | 24 | 25 |
| 4 | 42 | M | L | x | x | x |  | 17 | Y | N | 4 | 3.1 | 0.33 | 1.8 | 9.0 | 10 | 4.3 | 11 | 4.3 |
| 5 | 18 | F | L |  |  | x | x | 21 | Y | Y | 1 | 3.7 |  | 1.9 | 0.9 | 21 | 3.7 | 38 | 14 |
| 6 | 20 | F | R |  |  | x | x | 21 | Y | N | 3 | 2.7 | 1.49 | 2.6 | 3.0 | 9.1 | 12 | 15 | 17 |
| 7 | 22 | M | LR | x |  |  |  | 19 | Y | Y | 3 | 3.8 | 1.17 | 2.4 | 3.8 | 11 |  | 13 |  |
| 8 | 30 | F | R |  |  | x |  | 13 | Y | N | 5 | 5.2 | 1.26 | 6.5 | 14 | 19 |  | 20 |  |
| 9 | 43 | F | R | x | x |  |  | 12 | Y | N | 4 | 3.1 | 0.45 | 3.7 | 4.4 | 13 | 7.9 | 16 | 7.2 |
| 10 | 16 | M | R |  |  | x |  | 17 | Y | Y | 5 | 3.8 | 1.11 | 7.5 | 8.8 | 13 |  | 18 |  |
| 11 | 32 | F | R | x |  | x | x | 30 | Y | N | 3 | 5.1 | 3.43 | 8.4 | 2.8 | 7.3 | 7.7 | 12 | 12 |
| 12 | 36 | M | L |  |  |  | x | 25 | Y | Y | 4 | 5.2 | 1.48 | 11.6 | 6.5 |  | 5.5 |  | 7 |
| 13 | 21 | F | L | x | x | x |  | 14 | Y | N | 3 | 6 | 0.47 | 14.7 | 1.1 | 13 | 6.2 | 19 | 4.9 |
| 14 | 21 | F | R |  | x |  |  | 14 | Y | N | 8 | 3.7 | 1.1 | 16.9 | 9.3 |  | 11 |  | 15 |
| 15 | 29 | F | R |  | x |  | x | 17 | Y | N | 4 | 2.5 | 1.31 | 5.4 | 2.7 |  | 3.6 |  | 7.2 |
| 16 | 41 | F | R |  |  | x |  | 19 | Y | N | 3 | 4.6 | 0.72 | 7.4 | 4.4 | 9.4 |  | 11 |  |
| 17 | 24 | M | R | x |  |  |  | 21 | Y | N | 3 | 5.1 | 1.88 | 8.9 | 2.9 | 23 |  | 24 |  |
| 18 | 31 | F | R | x |  |  |  | 15 | Y | N | 6 | 6.1 | 1.18 | 24.2 | 3.8 | 7.8 |  | 10 |  |
| 19 | 21 | M | R | x | x |  |  | 15 | Y | Y | 4 | 2.8 | 0.72 | 5.4 | 5.8 | 12 | 2.6 | 19 | 6.1 |
| 20 | 19 | F | R |  |  | x |  | 21 | Y | Y | 3 | 4.9 | 2.53 | 5.2 | 6.0 | 11 |  | 12 |  |
| mean | 30 | F=13 | R=15 | N=11 | N=7 | N=9 | N=5 | 18 | N=20 | N=8 | 4 | 4.4 |  | 9.39 | 4.64 | 12 | 7 | 18 | 11 |

### Electrode localization

After implantation, electrodes were located by aligning post-implant CT to preoperative 3D T1-weighted structural MRI with ∼1mm^3^ voxel size (Dykstra et al., 2012), using 3D Slicer (RRID:SCR_005619). This allows visualization of individual contacts with respect to HC cross-sectional anatomy, which was interpreted in reference to published atlases (Duvernoy, 1988; Ding and Van Hoesen, 2015; Adler et al., 2018). While the spatial resolution of T1 MRI is limited, the body of HC has a highly regular anatomical orientation with respect to the temporal horn of the lateral ventricle. Specifically, when using a lateral to medial approach, the electrode first passes through the subtemporal white matter, then ventricle, then alveus, then field CA1, in the order of striata oriens, pyramidale, radiatum, and lacunosum, with considerable regularity (Duvernoy, 1988; Ding and Van Hoesen, 2015; Adler et al., 2018). With this anatomical guideline and the help of physiological activity (amplitude of sharp wave and ripple on unipolar recording), we assigned putative striatum pyramidale and radiatum to electrode contacts in HC (example shown in Fig. 1*A-B*). The assignment of depth contacts to anterior or posterior hippocampus (aHC/pHC) was made with the posterior limit of the uncal head as boundary (Poppenk et al., 2013; Ding and Van Hoesen, 2015).

**Figure 1.**
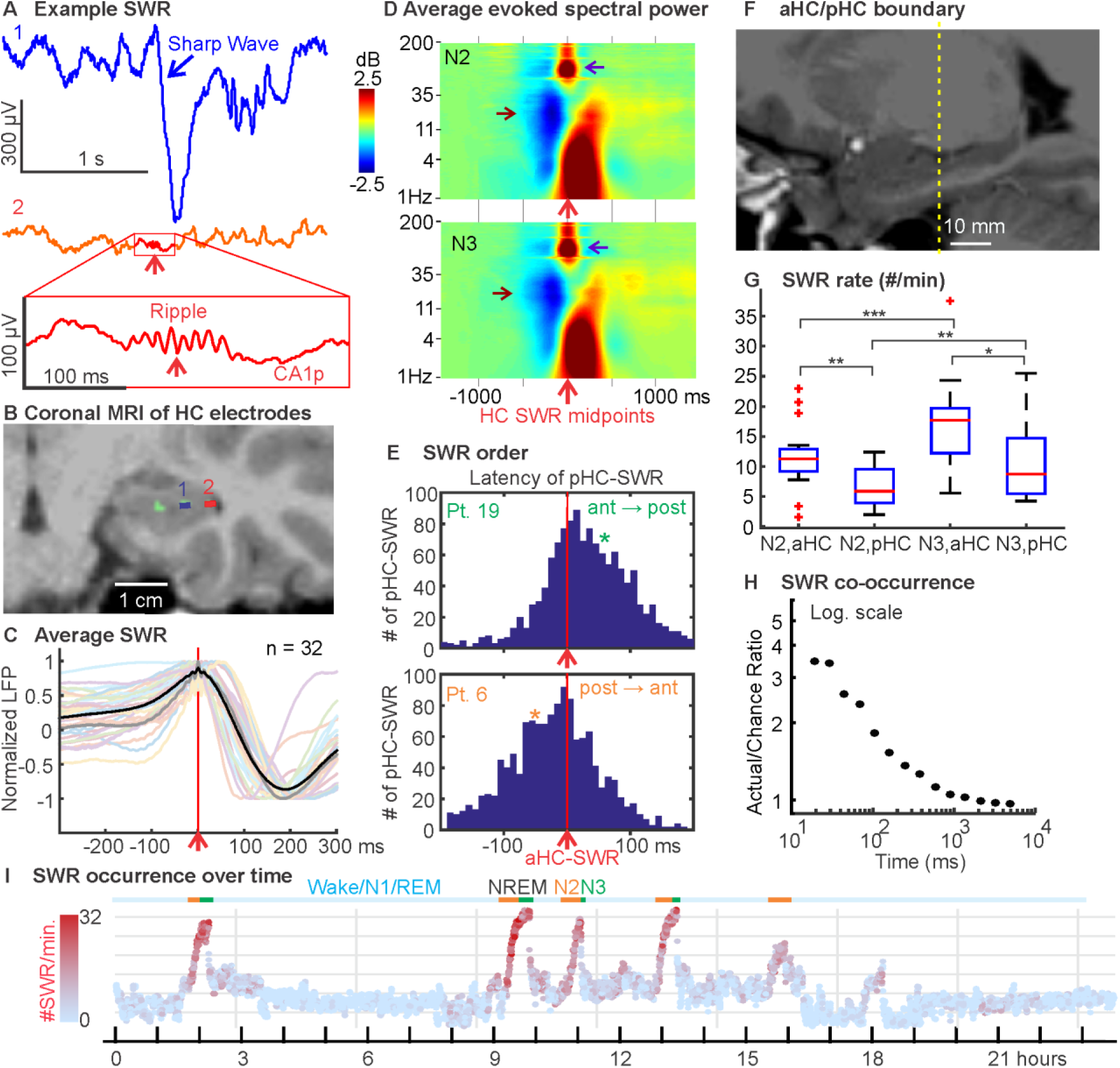
Characterization of human HC-SWR. ***A***. Single sweep of HC SEEG showing an example SWR, with the sharp wave prominent in putative CA1 stratum radiatum (1) and the ripple marked with rectangle on the LFP trace from putative CA1 stratum pyramidale (2, CA1p). ***B***. Coronal MRI of the contacts shown in A. ***C***. Overlaid average waveforms of SWR across hippocampal contacts from all patients, with the grand average across patients bolded in black. ***D***. Grand averages of all patients’ time-frequency plots of HC LFP triggered on ripple midpoints, showing typical frequency ranges of SWR (purple arrows), and broadband power decreases near SWR (brown arrows), from NREM stages N2 (top) and N3 (bottom). ***E***. In some patients (e.g., pt. 19), aHC SWR tend to precede pHC SWR, while the opposite was found in others (e.g., pt. 6). Detailed results and statistics can be found in Extended Data Figure 1-1. Each panel contains peri-stimulus time histograms over ±200 ms, with aHC-SWR as triggers (time 0, red vertical lines). Pt.: patient. Green/Orange stars mark the time bin closest to when an aHC/pHC SWR would reach pHC/aHC, respectively, given the previously estimated intrahippocampal SWR propagation speed of 0.35 m/s (Patel et al., 2013). ***F***. Sagittal MRI showing the boundary between aHC and pHC (marked in yellow), using the uncal apex criterion (Poppenk et al., 2013). ***G***. Box-and-whisker plot of SWR occurrence rates in different NREM stages across the HC longitudinal axis. Red plus signs mark outliers (>1.5 times the interquartile range). *: p < 0.05; **: p < 0.01; ***: p < 0.001. ***H***. For pairs of different HC sites (aHC/pHC or bilateral HC), SWR co-occurrence likelihood is low even at latencies <30 ms, and drops to chance in <1 s. Blue filled circles mark the ratios of actual over chance overlap proportions. ***I***. State plots showing the separation of NREM sleep in 24-hour LFP recording, using the first principal component derived from 24 vectors (one per cortical bipolar SEEG channel) of frequency power ratios (0.5-3 Hz over 0.5-16 Hz) (Gervasoni et al., 2004; Jiang et al., 2017). SWR rates are color coded with red intensity, and N2/N3 periods are marked with orange/green horizontal lines, respectively. Each data point covers 30 seconds, with 10-second overlap between two adjacent points.

Recordings were obtained from 32 HC contacts, 20 anterior (11 left) and 12 posterior (7 left). In 4 patients, HC recordings were bilateral (3 anterior and 1 posterior), and in 8 patients, ipsilateral anterior and posterior HC were both recorded. The distance of each hippocampal contact from the anterior limit of the hippocampal head was obtained in Freesurfer (RRID:SCR_001847). The CT-visible cortical contacts were compared to the MRI-visible cortical ribbon in order to identify pairs of contacts which are on the pial and white matter sides of the same cortical patch. These anatomically-selected pairs were confirmed physiologically by the presence of mainly polarity-inverted local field potentials in the referential recordings. Subsequent analysis was then confined to recordings between such bipolar transcortical pairs, yielding local field potential gradients. We have previously shown that referential recordings of sleep graphoelements can yield misleading localization, but that activity recorded by bipolar transcortical pairs is locally generated (Mak-McCully et al., 2015).

Electrode contacts were rejected from analysis if they were involved in the early stages of the seizure discharge, had frequent interictal activity or abnormal spontaneous local field potentials. From the total of 2844 contacts implanted in the 20 patients, 366 transcortical pairs (18.3±4.7 per patient) were accepted for further analysis. Polarity of the pairs was adjusted if necessary to “pial surface minus white matter” according to MRI localization, confirmed with decreased high gamma power during surface-negative downstates (see below).

Freesurfer (Dale et al., 1999; |Fischl et al., 1999a, 2004) was used to reconstruct from individual MRI scans the cortical pial and inflated surfaces, as well as automatic parcellation of the cortical surface into anatomical areas (Desikan et al., 2006), after a sulcal-gyral alignment process. In order to test for differences in HC-NC relationships between cortical locations, standard FreeSurfer ROIs were combined into 12 composite ROIs, and these were further combined across the two hemispheres in homologous ROIs (Extended Data Table 3-1; Fig. 5A). In order to test for HC-NC effects within vs between hemispheres, the ROIs were combined into Fronto-Central and Non-Frontal for each hemisphere (Table 3-1). An average surface for all 20 patients was then generated to serve as the basis of all 3D maps. While each cortical SEEG electrode contact’s location was obtained through direction correlation of CT and MRI as described earlier in this section, we obtained the cortical parcellation labels corresponding to each contact by morphing the Right-Anterior-Superior-oriented anatomical coordinates from individual surfaces to the average surface space (Fischl et al., 1999b). In addition, the nearest vertices on the average surface to the morphed coordinates would be identified for subsequent plotting of 2D projections onto the left lateral view. For the 2D projections only, to optimize visualization (i.e. minimize multiple contact overlap) while preserving anatomical fidelity, positions of contacts with significant HC-NC GE correlation were allowed to shift within a 5 mm radius. All visualizations were created with custom scripts in MATLAB 2016b.

### Data collection and preprocessing

Continuous recordings from SEEG depth electrodes were made with cable telemetry system (JE-120 amplifier with 128 or 256 channels, 0.016-3000 Hz bandpass, Neurofax EEG-1200, Nihon Kohden) across multiple nights (Table 1) over the course of clinical monitoring for spontaneous seizures, with 1000 Hz sampling rate. The SEEG referencing during recording was to a scalp electrode at Cz, while subsequent detection and analysis of cortical and hippocampal events were performed on bipolar derivations between adjacent contacts, to ensure signal focality (Mak-McCully et al., 2015), with only bipolar pairs that spanned the cortical ribbon and had no significant and/or persistent presence of pathological activity being kept for subsequent analysis. Out of the 2572 channels from the 20 patients we used for subsequent analysis, 480 were kept.

Recordings were anonymized and converted into the European Data Format (EDF). Subsequent data preprocessing was performed in MATLAB (RRID:SCR_001622); the Fieldtrip toolbox (Oostenveld et al., 2011) was used for bandpass filters, line noise removal, and visual inspection. Separation of patient NREM sleep/wake states from intracranial LFP alone was achieved by previously described methods utilizing clustering of first principal components of delta-to-spindle and delta-to-gamma power ratios across multiple LFP-derived signal vectors (Gervasoni et al., 2004; Jiang et al., 2017), with the addition that separation of N2 and N3 was empirically determined by the proportion of down-states that are also part of slow oscillations (at least 50% for N3 (Silber et al., 2007)), since isolated down-states in the form of K-complexes are predominantly found in stage 2 sleep (Cash et al., 2009).

The total NREM sleep durations vary across patients; while some difference is expected given intrinsic variability of normal human sleep duration (Carskadon and Dement, 2010), patients undergoing seizure monitoring experience additional disruption due to the hospital environment and clinical care. While most of our subsequent analyses utilize data from 20 patients over a total of 77 nights, in order to examine the potential effect of sleep disruption on our results, we identified 28 nights in 16 patients in which the percentages of NREM were comparable to (i.e. within 2 standard deviation of) normative data (Moraes et al., 2014) for separate analysis. In particular, we compared the occurrence rates of HC-SWR and NC-GE in NREM sleeps with normative N2 and N3 durations to their rates in other sleeps. We used linear mixed effect models (with patient ID being a random effect) to evaluate the significance of “normative” or “other” sleep categories on LFP event occurrence rates in N2 and N3, as detailed later in Methods under “Experimental design and statistical analysis: linear mixed effect models”. We found no significant effect of sleep categories in either NREM stage (p > 0.3964 and p > 0.0735 for all LFP event types in N2 and in N3, respectively).

### Hippocampal sharpwave-ripple (HC-SWR) selection

Previous studies have used a variety of approaches to identify HC-SWR, with regard to how they distinguish epileptiform interictal spikes from HC-SWR (or even if they do distinguish them). The literature is also inconsistent regarding the features used for detection, e.g. whether studies require that multiple oscillations are present in a ripple, or simply a peak in ripple-band power, or whether they require that the ripple occur at the peak of a HC sharpwave and followed by a slower wave of opposite polarity. In addition, the NREM sleep stage (N2 vs. N3) and the anatomical origin of human HC-SWR along the hippocampal longitudinal axis often go unreported (Bragin et al., 1999; Clemens et al., 2007; Axmacher et al., 2008; Brázdil et al., 2015; Staresina et al., 2015).

The HC-SWR we studied here were clearly distinguished from epileptiform activity by multiple criteria, possessed multiple peaks within the ripple and the ripple itself was located within a characteristic low frequency LFP. Specifically, HC LFP data were band-pass filtered between 60 and 120 Hz (6th order Butterworth IIR bandpass filter, zero-phase shift by forward and reverse filtering). This band was chosen based on examination of the time-frequency plots of visually-selected SWR in patients with no HC interictal spikes. Root-mean-square (RMS) over time of the filtered signal was calculated using a moving average of 20 ms, with the 80th percentile of RMS values for each HC channel being set as a heuristic cut-off. Whenever a channel’s signal exceeds this cut-off, a putative ripple event was detected. Adjacent putative ripple event indices less than 40 ms apart were merged, with the center of the new event chosen by the highest RMS value within the merged time window. Each putative ripple event was then evaluated based on the number of distinct peaks in the HC LFP signal (low-passed at 120 Hz) surrounding the event center; a 40 ms time bin was shifted (5 ms per shift) across ±50 ms, and at least one such time bin must include more than 3 peaks (the first and the last peak cannot be both less than 7 ms away from the edges of this time bin) for the putative ripple event to be considered for subsequent analyses. In addition, the distance between two consecutive ripple centers must exceed 40 ms.

### Rejection of epileptiform activity

Since RMS peaks may arise from artifacts or epileptiform activity, 2000 ms of hippocampal LFP data centered on each ripple event undergoes 1-D wavelet (Haar and Daubechies 1-5) decompositions for the detection and removal of sharp transients (i.e. signal discontinuities). For each wavelet decomposition, putative HC-SWR were marked in ∼10 min long NREM sleep period (marking ended sooner if 400 putative HC-SWR had been found). A scale threshold was then established via iterative optimization for the best separation between hand-marked true ripple events and interictal events in the same NREM sleep period. Each putative sharp transient was then rejected only if the 200 Hz highpassed data at that point exceeds an adaptive threshold (Bragin et al., 1999) of 3—or another number that allows best separation in agreement with visual inspection by human expert (between 0.5 and 5, and ∼2 on average)—standard deviations above the mean for the 10-second data preceding the transient.

To identify ripples coupled to sharpwaves, we created patient-specific average templates (−100 ms to +300 ms around ripple center) from hand-marked exemplars (100-400 HC-SWR in NREM per patient) that resemble previously described primate sharpwave-ripples with biphasic LFP deflections (Skaggs et al., 2007; Ramirez-Villegas et al., 2015). We then evaluated whether each ripple qualifies as a SWR based on its accompanying biphasic waveform with the following two criteria: 1, the similarity of the peri-ripple LFP to the average template as quantified by the dot product between the template and the peri-ripple LFP (−100 ms to +300 ms). For each template-LFP pair, the similarity threshold was chosen so that it would reject at least 95% of the hand-marked ripples with no sharpwave. 2, the absolute difference between the LFP value at the ripple center and at the maximum/minimum value between +100 ms and +250 ms after ripple center was computed for each ripple; this difference must exceed 10% (or another value which excludes 95% of hand-marked ripples with no sharpwave) of the difference distribution created from hand-marked SWR. In addition, the distance between two consecutive SWR centers must exceed 200 ms. Time-frequency plots of HC LFP centered on detected SWR were then created in MATLAB with EEGLAB toolbox (Delorme and Makeig, 2004), each trial covering ±1500 ms around SWR and with −2000 ms to −1500 ms as baseline (masked for significance at alpha = 0.05, 200 permutations).

Hippocampi involved early in the seizure discharge were rejected from analysis. Some of the remaining had no interictal spikes (IIS), whereas others did at a low level. We tested if the HC-SWR to NC-GE associations were affected by a low level of IIS with linear mixed effect models with the percentage of HC-NC channel pairs showing significant association as the response variable, the IIS-free/not-free as the predictor variable, and the subject-wise variability as random effect. We found no significant effect of IIS freedom on HC-NC association percentages (p = 0.9197). While this result could not preclude HC-NC connectivity being altered by pathology, it does not provide evidence for such alterations.

### Neocortical graphoelement (NC-GE) selection

Automatic cortical theta burst (TB) detection was performed as previously described (Gonzalez et al., 2018): a zero-phase shift 5-9 Hz bandpass was applied to the LFP data, and a channel-wise cutoff set at mean +3 s.d. was applied to each channel’s Hilbert envelope (smoothed with 300 ms Gaussian kernel) to identify local maxima that would be “peaks” of putative TBs, and make start/stop edge cut-offs at mean +1 s.d.. The resulting TBs were only accepted for further analyses if their durations fall between 400 ms and 1000 ms, with their frequency between 5 and 9 Hz as calculated from the number-of-zero-crossings.

Automatic cortical spindle (SS) detection was performed using previously published methods for SEEG spindle identification (Mak-McCully et al., 2017): a zero-phase shift 10–16 Hz bandpass was applied to the LFP data, and the analytic amplitude of the bandpassed signal was convolved with an average Tukey window of 600 ms. A channel-wise cutoff set at mean +2 s.d. was applied to each channel’s convolved bandpass envelope to identify local maxima that would be “peaks” of putative spindles. A 400 ms Tukey window was then applied to the previously computed Hilbert amplitude to define the onset and offset of putative SS; the maximum FFT amplitude of the putative SS signal at the SS peak was used to set edge thresholds (50 % of peak amplitude). Finally, the putative SS segments were evaluated with four different criteria for rejection: 1, the spindle duration needs to be longer than 400 ms; 2, the spectral power outside the spindle frequency band on the lower end (4-8 Hz) or the higher end (18-30 Hz) should not exceed 14 dB; 3, each spindle should consist of at least 3 unique oscillation peaks in the 10-16 Hz bandpass; 4, the time between successive zero-crossings should be within the range of 40-100 ms.

Downstates (DS) and upstates (US) were identified as follows: for each cortical LFP signal, a zero-phase shift application of eighth order Butterworth filter from 0.1 to 4 Hz was applied.

Consecutive zero crossings of opposite slope separated by 250 to 3000 ms were then selected as delineating putative graphoelements. For each putative graphoelement, the amplitude peak between zero crossing was computed; only the top 10% of peaks were retained. The polarity of each signal was inverted if necessary to assure that NC-DS were negative, as confirmed with a high gamma (70-190 Hz) power decrease exceeding 1 dB within ±250 ms of the negative NC-DS peaks.

### Experimental design and statistical analysis: linear mixed effect models for comparisons among GE occurrence rates and HC-NC association percentages

As mentioned previously in the Methods section under “Data collection and preprocessing”, in order to account for the possibility that patient-wise variations in NREM sleep durations beyond normative range would unduly influence our results, we identified 28 sleeps from 16 patients with normative N2 and N3 sleep durations. We then compared the occurrence rates of SWR and NC-GE in this subset of sleeps to the other 49 sleeps from 20 patients using linear mixed effect models (implemented in MATLAB as part of the Statistics and Machine Learning Toolbox). Specifically, for each NC-GE type (and HC-SWR) in a given NREM stage, we created three separate models: a baseline model where GE occurrence rates are predicted by whether a given sleep is in the “normative” or the “other” category (GE Rate ∼ 1 + Sleep Category), a model including patient random effect as a random intercept term (GE Rate ∼ 1 + Sleep Category + (1 | Patient ID)), and a model with patient random effect as both random intercept and slope terms (GE Rate ∼ 1 + Sleep Category + (Sleep Category | Patient ID)). Comparisons between model fits were conducted with likelihood ratio tests, with the base model being the best fit for all NC-GE types and NREM stages.

Similarly, we compared the occurrence rates of HC-SWR from different NREM stages (N2 vs. N3) and from different sources (aHC vs. pHC). Our base model has HC-SWR rates as the response variable, with AP (aHC/pHC) and Stage (N2/N3) as categorical predictor variables. The full model includes interaction between predictor variables and all possible random slope/intercept terms (SWR Rate ∼ 1 + AP*Stage + (AP | Patient ID) + (Stage | Patient ID) + (1 | Patient ID)), with reduced models’ fits being compared to the full model fit via likelihood ratio tests. The best-fit model turned out to be without interaction between predictor variables (SWR Rate ∼ 1 + AP + Stage + (1 | Patient ID)).

We also used linear mixed effect models to compare the percentages of HC-NC channel pairs that show significant NREM HC-SWR to NC-GE association (as evaluated in the Methods subsection “Co-occurrence of HC-SWR and NC-GE” below) in IIS-free versus other patients. The percentages serve as the response variable, while the “IIS-free or not” status for each patient serves as a categorical predictor variable, with patient-wise random effect. Comparisons between model fits between the baseline model (HC-NC association ∼ Patient Category) and the model with random intercept (HC-NC association ∼ 1+ Patient Category + (1 | Patient ID)) were conducted with likelihood ratio tests, with the baseline model being the better fit.

### Experimental design and statistical analysis: Time-frequency analysis

For time-frequency analysis in Figure 1, spectral content of the LFP data from hippocampal channels were evaluated using EEGLAB (RRID:SCR_007292) routines that applied wavelet transforms (Delorme and Makeig, 2004). Spectral power was calculated over 1 to 200 Hz across each individual time period (“trial”) centered around HC-SWR ripple centers found in NREM by convoluting each trial’s signal with complex Morlet wavelets, and averages were then taken across trials. The resulting time-frequency matrices were normalized with respect to the mean power at each chosen frequency and masked with two-tailed bootstrap significance at α = 0.05, with the pre-trigger times (−2000 ms to −1500 ms) as baseline.

### Experimental design and statistical analysis: SWR co-occurrence over multiple HC sites

We evaluated in patients with multiple hippocampal recording sites the extent to which SWR overlap (i.e. fall within 50 ms of each other). This was tested both for two sites within the same hippocampus along the longitudinal axis (anterior and posterior), or in the two hemispheres (left and right, in the same longitudinal position). To test whether such overlaps differ from expected values due to chance alone (with variable overlap criteria being the time window size—ranging from 25 to 5000 ms—within which SWR from multiple HC sites co-occur), we estimated the co-occurrence rate for random SWR times, with the proviso that the total number of randomly-occurring SWR in each 5 min interval matches that actually observed. One-tailed 2-sample t-tests were then conducted between actual co-occurrences versus expected, with one pair of observations from each patient (N=8 for anterior vs posterior, and N=4 for left vs right). To test whether anterior led posterior HC-SWR in each patient we conducted a binomial test for the SWR pairs occurring in both sites within 200 ms, against the null hypothesis of equal numbers from anterior vs posterior leads.

### Experimental design and statistical analysis: Co-occurrence of HC-SWR and NC-GE

Statistical tests that involve multiple comparisons in this and subsequent Methods subsections all have α = 0.05 post-FDR correction (Benjamini and Hochberg, 1995). The FDR correction procedure was implemented as follows: for a total of N hypotheses, each with corresponding p-values P*_i_* (*i* ∈ N), which are sorted in ascending order to identify P*_k_* (*k* being the largest *i* for which P*_i_* ≤ (*i* / N) * α), all hypothesis with p-values less than or equal to P*_k_* would be rejected.

Peri-stimulus time histograms were constructed for each HC-NC channel pair, separately for each graphoelement (GE; TB, SS, DS, and US), and for each sleep-stage (N2 or N3), each histogram comprising the occurrence times of a given NC-GE during the ±2 s interval surrounding midpoints of HC-SWR (see Fig. 2 for examples). Altogether, there were 598 unique HC-NC pairs (458 ipsilateral, 140 contralateral), and 8 histograms for each pair (one for each sleep stage, times each of the 4 GEs), for a total of 4784 histograms. Overall, each of these histograms plots the occurrence times of 3048±4221 NC-GEs (+2.84 skewness) with respect to 2080±2502 HC-SWR (+2.1 skewness); a total of 14.6 million NC-GE events and 9.95 million HC-SWR were used for histogram construction. The significance of peaks and troughs in each histogram was tested by comparing them to distributions derived from histograms constructed under the null hypothesis of no relationship between the NC-GE and HC-SWR using the following randomization procedure. Null-hypothesis histograms (N=1000) were constructed of NC-GE occurrences relative to a number of random times equal to the number of HC-SWR. For each 200 ms time bin with 100 ms overlap comprising the 4-second trial duration, the actual counts are then compared to the distribution under the null hypothesis, followed by FDR correction.

**Figure 2.**
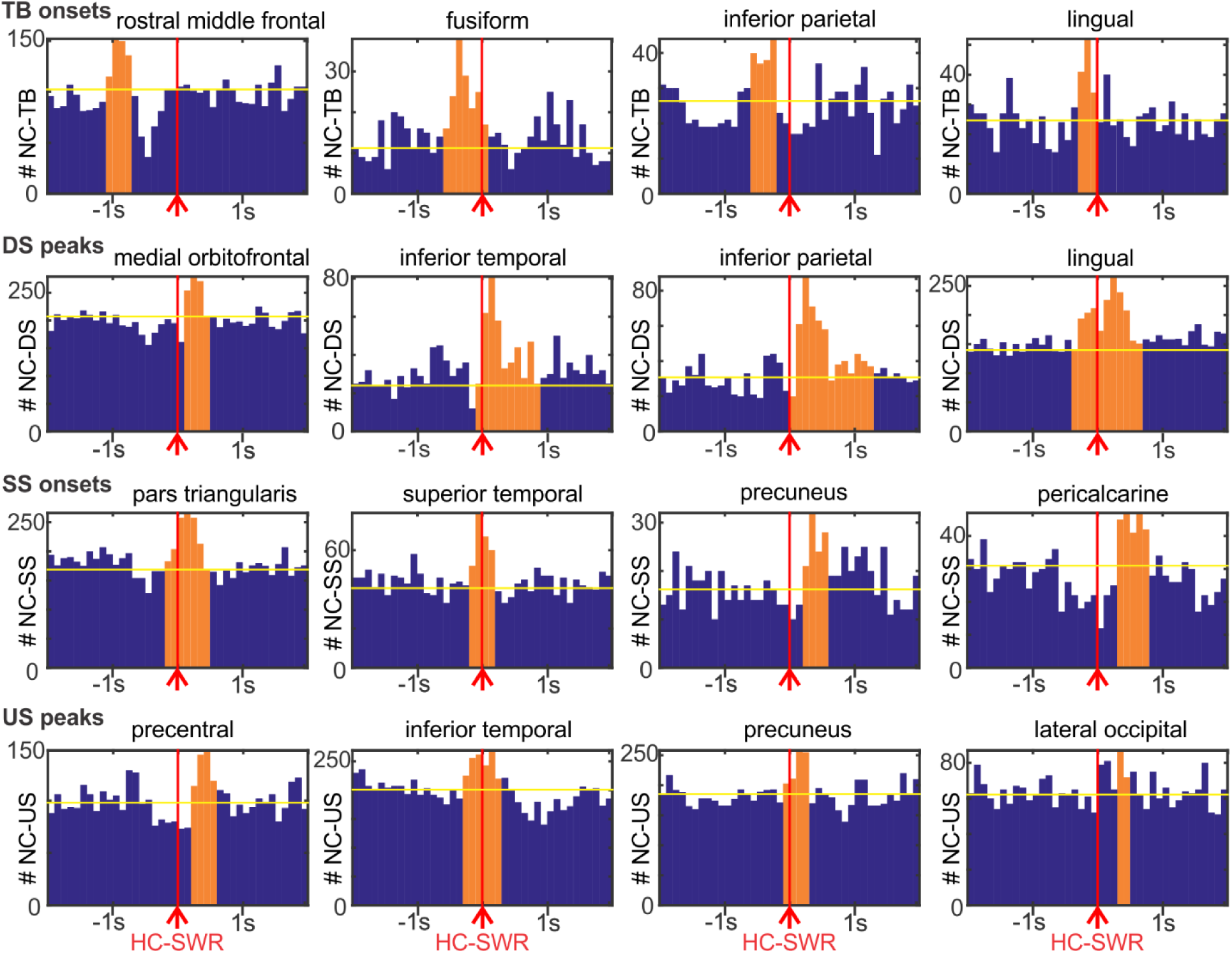
Neocortical graphoelements (NC-GE) in relation to HC-SWR, for example channel-pairs. Each row of histograms is for a different type of GE: theta burst onsets (TB), downstate peaks (DS), spindle onsets (SS), and upstate peaks (US). The four columns of plots show example histograms with significant temporal correlations between HC-SWR and NC-GE from the frontal, temporal, parietal, and occipital lobes, respectively. Red vertical lines indicate trigger (SWR) location for the peri-stimulus histograms. Orange bars indicate the time ranges with both peak NC-GE occurrence rate and significant correlation. Yellow horizontal lines show the mean baseline established via averaging randomized control distributions. A summary of the numbers of events across histograms can be found in Extended Data Figure 2-1.

The latencies of the largest significant peaks identified in the individual histograms constructed as described above, were used to create summary histograms for each NC-GE, sleep stage, and HC-SWR origin in anterior versus posterior HC. These are plotted in Fig. 3 and Supplementary Figure 3-1, and tabulated in Table 2. Each of these 16 histograms of histogram peaks (N2/N3 × aHC/pHC × 4 GEs) summarizes the significant latencies from 108±54 HC-NC channel pair histograms, comprising a total of 4.03 × 10^5^±3.78 × 10^5^ HC-NC events (range 2.44 × 10^4^-1.35 × 10^6^). To test if, overall, a given type of cortical graphoelement significantly precedes or follows HC-SWR, two-tailed binomial tests with chance probability of 0.5 were performed on the number of channel-pairs with peak latencies in the 500 ms before vs the 500 ms after the reference HC-SWR. Since certain graphoelements, such as cortical theta, tend to yield peak latencies centered around zero, we also tested with Kolmogorov-Smirnov tests whether, overall, for a given cortical-GE, the distribution of peak latencies significantly related to HC-SWR differs from chance.

**Table 2.** Relation of NC-GE to HC-SWR occurrences. Separate values and statistical significance tests are shown for NREM sleep stages N2 vs. N3, for anterior vs. posterior HC, and for different graphoelements (TB-theta bursts; DS-downstates; SS-sleep spindles; US-upstates). Tests indicate if there was a significant association between the times of occurrence of the SWR and GE in the overall histogram of significant histogram peaks (shown in Figs. 3 and 3-1) (fourth column); if the GE occurred significantly before the SWR (negative numbers) or before, when measuring the peak latency as the mean value across cortico-HC pairs (fifth column), or as the peak of an extreme value distribution fitted over the ±500 ms histogram-of-histograms in Figs. 3 and 3-1 (sixth column). The seventh column shows if there is a significant difference in the number of GE occurring 500 ms before vs. 500 ms after the SWR. The eighth column contains results from the same test as the seventh column, but applied to data with N2/N3/aHC/pHC combined. **BOLD** indicates significant values; *italics* indicate trends. For columns 4, 7, and 8, extremely small (< 0.0001) p-values are represented by 0 for clarity. Ant: anterior; Post: posterior.

| GE Type | Sleep Stage | HC Longitudinal Axis | SWR v. GE related (KS test p-value) | Peak Latency (ms), Mean | Peak Latency (ms), DistPeak | Non-zero Latency, binomial p-value | Non-zero Latency, binomial p-value, overall |
| --- | --- | --- | --- | --- | --- | --- | --- |
| TB onset | N2 | Ant | **0.023** | -92 | -104 | 0.238 | **0.0011** |
|  |  | Post | **0** | -45 | -123 | 0.065 |  |
|  | N3 | Ant | **0.023** | 0 | -95 | 0.508 |  |
|  |  | Post | **0.003** | -62 | -209 | 0.052 |  |
| DS peak | N2 | Ant | **0** | 0 | 94 | 0.228 | **0** |
|  |  | Post | **0** | **186** | **356** | **0** |  |
|  | N3 | Ant | **0.002** | 35 | 236 | 0.091 |  |
|  |  | Post | **0** | **128** | **224** | **0** |  |
| SS onset | N2 | Ant | 0.115 | **150** | **179** | **0.009** | **0.0002** |
|  |  | Post | **0.02** | **187** | **309** | **0.001** |  |
|  | N3 | Ant | 0.452 | -32 | -229 | 0.481 |  |
|  |  | Post | 0.073 | 238 | 342 | 0.092 |  |
| US peak | N2 | Ant | **0** | **126** | **216** | **0** | **0** |
|  |  | Post | **0** | **164** | **279** | **0** |  |
|  | N3 | Ant | **0** | **76** | **134** | **0** |  |
|  |  | Post | **0** | **81** | **85** | **0.013** |  |

**Figure 3.**
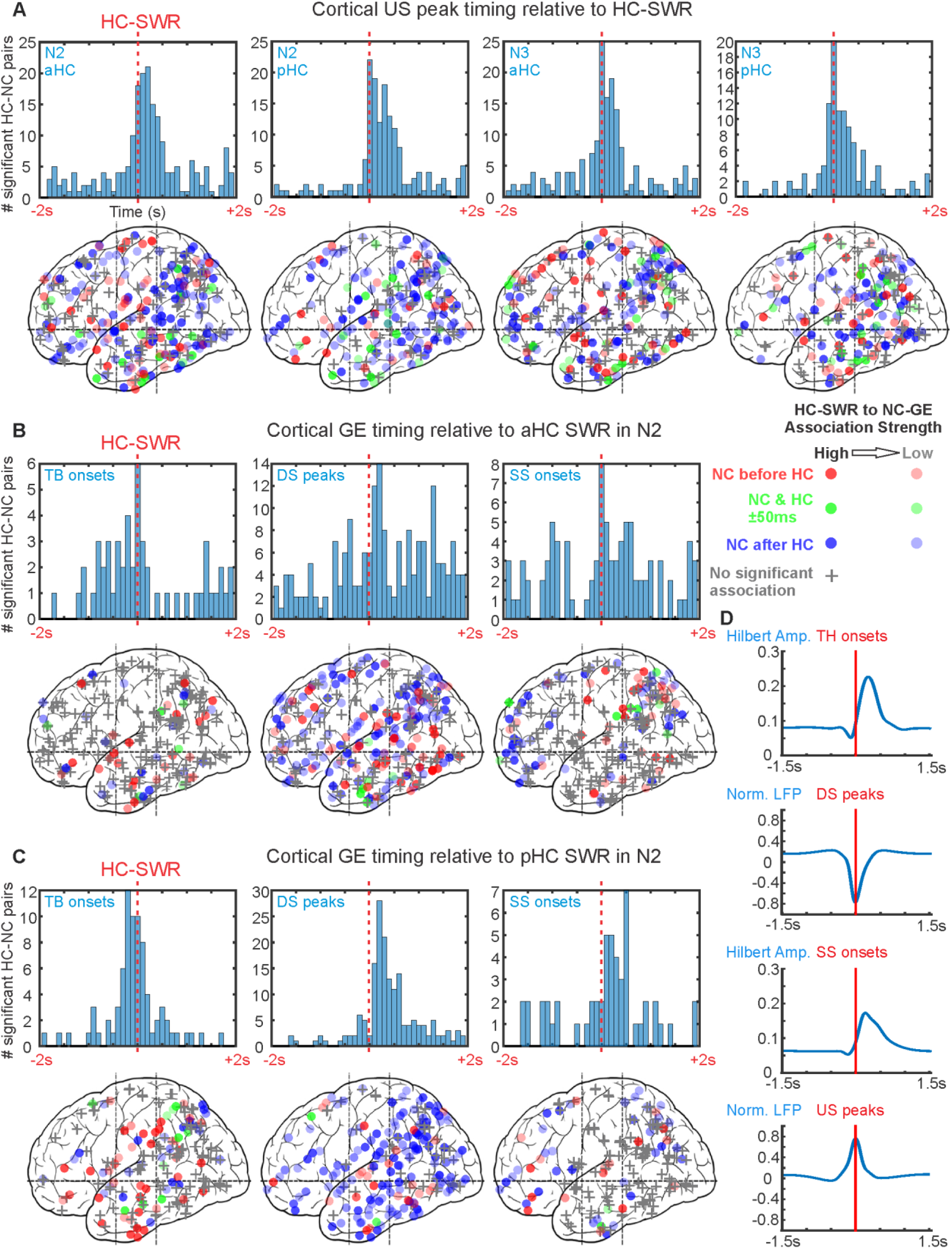
NC-GE tend to co-occur with HC-SWR at consistent latencies. ***A-C***. *Top*: Histograms of peak latencies for HC-NC channel pairs with significant temporal correlations between HC-SWR and NC-GE. Each count is the peak latency of a particular HC-NC channel pair (i.e., each histogram in Fig. 2 would contribute one count to one of the histograms in this figure). *Bottom*: maps of peak latency between SWR and GE for cortical channels. Circles indicate where GE-SWR relationships are significant, while plus signs represent non-significant channels. The intensity of each circle corresponds to the strength of GE-SWR coupling estimated (as in Fig. 2), and the color of each circle indicates peak latency: red for GE before SWR, blue for GE after SWR, and green for GE co-occurring with SWR (i.e. within 50 ms of each other). Both hemispheres, and medial and lateral cortical sites, are superimposed in each plot. ***A***. NC-US tend to follow both aHC and pHC SWR in both N2 and N3. ***B***. NC-TB tend to precede pHC SWR, while other NC-GEs tend to follow. ***C***. NC-TB tend to precede aHC-SWR, while NC-SS tend to follow. ***B-C.*** Stage N2 NREM sleep. Stage N3 results are shown in Extended Data Figure 3-1, and down-to-up state transition results are shown in Figure 3-2. Statistics from further analyses on the non-homogeneity of spatial distributions for NC sites significantly coupled to HC-SWR can be found in Figures 3-3, 3-4, and 3-5. ***D***. average traces of NC-GE Hilbert amplitude (for TB/SS) and LFP (for DS/US) centered on TB/SS onsets and DS/US peaks, showing the durations of each NC-GE type.

### Experimental design and statistical analysis: Differences in anatomical patterns of NC-GE relationship to HC-SWR between anterior vs posterior and left vs right HC

For every HC site across all patients, we tested the following: 1. whether the anatomical distribution of NC-GE coupling to HC-SWR across NC sites was different from chance for a given HC site. Subsequently, we tested the following: 2. whether the significantly-different-from chance distributions of NC sites from 1 would also differ between HC site pairs. An HC site pair could be either aHC and pHC in the same hippocampus, or left HC and right HC (either both aHC or both pHC) in the same patient; 3. whether any difference seen in 2 could be seen on the basis of different NC-GE types.

For 1, we performed a chi-square test of homogeneity for each HC site with regard to each NC-GE type, i.e. for a given patient with multiple NC sites, a 2 × N contingency table was made for each NC site, N being the number of NC sites, one column of table containing the number of NC-GE overlapping with (i.e. falling within 500 ms of) HC-SWR for each NC, the other containing the number of NC-GE that do not. A significant chi-square (post FDR-correction) would therefore indicate a significant non-random distribution for that particular HC site and NC-GE type.

For 2, we performed Wilcoxon signed rank tests on the data obtained in 1 to evaluate whether for a given NC-GE type (for which 1 yielded significance for both members of an HC site pair), the proportions of NC-GE from significantly coupled NC sites that overlapped with aHC/left HC-SWR differ from the proportions for pHC/right HC-SWR. Each test required that both members of a HC site pair must have significant chi-square test result from 1.

For 3, we repeated the procedure in 2, but instead of comparing proportions between HC site pairs, we compared between NC-GE types for each HC site. Each test required that for a given HC site, both NC-GE types were required to have significant chi-square test result from 1. Analyses 1-3 were performed with N2 and N3 combined.

### Experimental design and statistical analysis: Modulation of HC-SWR with GE-DS co-occurrence by number of DS

HC-SWR were observed to commonly be surrounded by broadband decreased spectral power, suggesting a possible relationship to NC-DS. To test whether multiple NC-DS are more likely than single NC-DS to co-occur with HC-SWR, we generated the baseline occurrence rate of single DS and multiple DS (averages derived from 200 permutations with pseudo-SWR) within ±500 ms of SWR for all HC contacts in all patients. We then obtained the ratios between the numbers of actual single DS / multiple DS and the baselines, followed by paired t-tests against the null hypothesis of equal mean for single DS and multiple DS ratios across patients. In order to account for patient-wise variability, we also built linear mixed-effect models with random intercept and/or slope for patient ID, SWR-DS co-occurrence rate as response variable, and single vs. multi-DS as categorical predictor variable. The best-fit model has random intercept (SWR-DS rate ∼ 1 + DS type + (1 | Patient ID)).

### Experimental design and statistical analysis: Canonical sequences of NC-GE

Previous studies have identified canonical sequences of different NC-GE in human NREM SEEG, specifically TB-DS-SS (Mak-McCully et al., 2017; Gonzalez et al., 2018). Accordingly, we identified triplet sequences (involving one TB, one DS, and one SS from NC) near HC-SWR, with the temporal limitation that TB start, SS start, and DS peak all must fall within −1500 ms and +1000 ms of an HC-SWR to yield a GE triplet. To test whether previously observed canonical order would be preferred for these triplets near HC-SWR, we tallied the number of triplets occurring in each of the six possible orders and performed binomial tests against chance (1/6), and computed the rates of occurrence for all possible triplets in NREM. Since SS starts and US peaks tend to overlap in their latency distributions with regard to HC-SWR, we repeated this analysis with SS and US combined (so a typical sequence would be TB-DS-SS/US).

### Experimental design and statistical analysis: Modulation of HC-SWR co-occurrence with NC-GE by sleep stage (N2 vs N3), cortical site (anatomical ROI), HC site (anterior vs posterior), NC-GE type, and laterality (HC-NC ipsilateral or contralateral to each other)

We further explored the spatial distribution of the HC-NC relationship by tallying the proportion of significant HC-NC channel pairs across different NC regions, with respect to different NC-GE types and different HC-SWR sources (Table 3). To evaluate the statistical significance of this distribution, we performed a 4-way ANOVA, comparing the main effects of GE type (TB, SS, DS, US), NREM stage (N2 vs. N3), a/pHC origin of HC-SWR, and NC ROIs (coverage listed in Table 3 and Table 3-1), as well as the 2-way and 3-way interaction effects. Similarly, given the lateralization of human hippocampal function, we were interested in examining whether ipsilateral HC-NC channel pairs would show different relationships from contralateral pairs across GE types, NREM stages, or NC ROIs. Due to a sparse representation of contralateral channel pairs for individual ROIs, we calculated proportions for larger regions by combining both HC and NC sites: aHC and pHC channels were combined into a single HC category, and the 10 NC ROIs used in previous analysis were collapsed into two: “fronto-central” and “non-frontal” (Table 3-1). We then computed for each ROI the proportion of NC channels significantly coupled to HC-SWR (Table 5), and performed 4-way ANOVA as previously described, with the ipsilateral/contralateral factor replacing the a/pHC factor. All post-hoc analyses for both ANOVAs were performed with Tukey’s range test. For each ANOVA, we checked the normality assumption by conducting the Lilliefors test on the residuals.

**Table 3.** Numbers of HC-NC channel pairs by NC regions of interest (ROIs) and proportions of HC-NC channel pairs with significant NC-GE/HC-SWR relationships across NC ROIs. Ch. pairs: HC-NC channel pairs. Details regarding the Freesurfer parcellations included in each ROI can be found in Extended Data Table 3-1.

|  |  |  | Proportion of channels showing significant NC-GE co-occurrence with HC-SWR |  |  |  |  |  |  |  |  |  |  |  |  |  |  |  |
| --- | --- | --- | --- | --- | --- | --- | --- | --- | --- | --- | --- | --- | --- | --- | --- | --- | --- | --- |
| GE Type | Total# Ch. pairs |  | TB |  |  |  | DS |  |  |  | SS |  |  |  | US |  |  |  |
|  |  |  | N2 |  | N3 |  | N2 |  | N3 |  | N2 |  | N3 |  | N2 |  | N3 |  |
| ROI | aHC | pHC | aHC | pHC | aHC | pHC | aHC | pHC | aHC | pHC | aHC | pHC | aHC | pHC | aHC | pHC | aHC | pHC |
| Lateral occipital | 15 | 15 | 0.067 | 0.067 | 0.000 | 0.133 | 0.200 | 0.467 | 0.200 | 0.400 | 0.000 | 0.067 | 0.267 | 0.000 | 0.267 | 0.600 | 0.200 | 0.400 |
| Medial temporo-occipital | 31 | 26 | 0.194 | 0.077 | 0.032 | 0.192 | 0.516 | 0.808 | 0.387 | 0.423 | 0.129 | 0.231 | 0.097 | 0.192 | 0.645 | 0.615 | 0.419 | 0.308 |
| Lateral parietal | 74 | 51 | 0.081 | 0.412 | 0.041 | 0.137 | 0.392 | 0.686 | 0.365 | 0.529 | 0.203 | 0.157 | 0.203 | 0.059 | 0.419 | 0.765 | 0.446 | 0.431 |
| Medial occipito-parietal | 34 | 27 | 0.029 | 0.259 | 0.059 | 0.111 | 0.500 | 0.667 | 0.529 | 0.519 | 0.529 | 0.185 | 0.265 | 0.296 | 0.412 | 0.630 | 0.500 | 0.481 |
| Lateral temporal | 86 | 54 | 0.186 | 0.407 | 0.070 | 0.204 | 0.558 | 0.889 | 0.372 | 0.537 | 0.140 | 0.222 | 0.047 | 0.037 | 0.640 | 0.685 | 0.407 | 0.556 |
| Insula | 11 | 6 | 0.455 | 0.667 | 0.091 | 0.167 | 0.909 | 0.667 | 0.364 | 0.167 | 0.000 | 0.000 | 0.000 | 0.000 | 0.727 | 0.667 | 0.545 | 0.667 |
| Paracentral | 29 | 16 | 0.103 | 0.250 | 0.172 | 0.063 | 0.724 | 0.625 | 0.517 | 0.625 | 0.276 | 0.125 | 0.034 | 0.000 | 0.828 | 0.625 | 0.517 | 0.688 |
| Orbitofrontal | 23 | 6 | 0.217 | 0.667 | 0.130 | 0.333 | 0.565 | 1.000 | 0.478 | 0.167 | 0.435 | 1.000 | 0.174 | 0.167 | 0.478 | 1.000 | 0.348 | 0.167 |
| Prefrontal | 49 | 14 | 0.061 | 0.286 | 0.082 | 0.429 | 0.551 | 0.857 | 0.469 | 0.786 | 0.367 | 0.714 | 0.204 | 0.286 | 0.429 | 0.786 | 0.510 | 0.500 |
| Cingulate | 13 | 6 | 0.231 | 0.667 | 0.385 | 0.667 | 0.462 | 0.833 | 0.538 | 0.833 | 0.231 | 0.167 | 0.308 | 0.167 | 0.615 | 0.667 | 0.692 | 1.000 |
| Total# Ch. pairs across NC | 365 | 221 |  |  |  |  |  |  |  |  |  |  |  |  |  |  |  |  |
|  | Average proportion |  | 0.162 | 0.376 | 0.106 | 0.244 | 0.538 | 0.750 | 0.422 | 0.499 | 0.231 | 0.287 | 0.160 | 0.120 | 0.546 | 0.704 | 0.459 | 0.520 |
|  | Standard deviation |  | 0.119 | 0.219 | 0.104 | 0.175 | 0.178 | 0.147 | 0.100 | 0.212 | 0.167 | 0.300 | 0.103 | 0.110 | 0.163 | 0.115 | 0.124 | 0.218 |

### Code Accessibility

All custom scripts would be available upon request by contacting the corresponding author and would be delivered through the UCSD RDL-share service.

## Results

### Characterization of human hippocampal sharpwave-ripples (HC-SWR)

We identified human HC-SWR from intracranial recordings (Fig. 1*A-B*), characterized their density and spectral traits, and determined if their characteristics change between NREM stages N2 and N3, or between anterior and posterior HC sites (as defined by the uncal head boundary; see Fig. 1*F*). Morphologically normal HC-SWR were isolated from >24 h continuous recordings in 20 stereoelectroencephalography (SEEG) patients with intractable focal epilepsy (Fig. 1*C*), with anterior (in 17 patients) and/or posterior HC contacts (in 11 patients), and related to sleep graphoelements (GE) in 12-30 NC bipolar recording channels per patient.

We found clear differences in the occurrence rates of HC-SWR in different NREM stages and in aHC vs. pHC (Table 1, Fig. 1*G*). Specifically, for aHC, the mean SWR rate was 11.59/min in N2, and significantly greater at 17.01/min in N3 (two-tailed paired t-test, p = 0.0001); for pHC, the mean SWR rate was 6.75/min in N2, and significantly greater at 10.71/min in N3 (two-tailed paired t-test, p = 0.0083). Comparing aHC SWR rate to pHC SWR rate under different NREM stages, in turn, also yielded significant differences, with aHC consistently producing more SWR than pHC (two-tailed t-tests, p = 0.0038 for N2, p = 0.018 for N3). In contrast to NREM, SWR density was low in waking (Fig. 1*I*) for aHC (1.77/min) and pHC (2.16/min), which did not differ significantly (p = 0.5433, two-tailed two-sample t-test).

Since the aHC/pHC and N2/N3 differences in SWR rates above might be due to patient-wise variability, we also built linear mixed-effect models with SWR rate as the response variable, being predicted by patient identity (random effect), SWR source (aHC/pHC, categorical variable), and NREM stage (N2/N3, categorical variable). The best-fit model has random intercept due to patient identity, while both SWR source and NREM stage still had significant effects on SWR rate (p = 6.552 × 10^−5^ and p = 0.0001, respectively). Thus, the aHC and N3 preferences of SWR occurrence cannot be solely accounted for by sampling error, and may instead be physiological in origin.

Within the HC, SWR were surrounded by locally-decreased broadband spectral power (Fig. 1*D*). The post-SWR decrease appears more prominently in N3 and limited to >60 Hz; previous studies in rodents have shown that the delta-wave (which typically follows the sharpwave) is accompanied by hyperpolarization of HC pyramidal cells (English et al., 2014). However, here we observed an additional, stronger (0.55-1 dB below baseline), longer and broader (<120 Hz) broadband decrease in spectral power which commonly began 500-1000 ms prior to the ripple (statistically significant at α = 0.05; see “Experimental design and statistical analysis: Time-frequency analysis” in Methods).

### Co-occurrence of HC-SWR across different hippocampal locations

Although initial descriptions of SWR in rodents emphasized their co-occurrence throughout the entire extent of both hippocampi (Buzsaki et al., 1992), more detailed studies showed that isolated SWR could also occur (Patel et al., 2013), suggesting that different populations of SWR may interact with neocortical events in disparate regions. In our data, for patients (n = 8) with both aHC and pHC contacts in the same HC (Fig. 1*E*, Fig. 1-1*A*), the overlap (i.e. co-occurrence within 50 ms) between aHC and pHC SWR was 5.6±3.0% in N2, and 7.3±5.9% in N3, compared to chance levels of 1.0±0.54% in N2 and 1.6±1.0% in N3 (Fig. 1-1*B*). In both N2 and N3, the actual mean overlap percentages were significantly greater than the mean overlap percentages derived from chance (p = 0.0018 and p = 0.011, one-tailed paired t-tests). By varying the co-occurrence criterion range between 25 ms and 5000 ms, we observed greater-than-chance co-occurrence rates for aHC and pHC SWR initially, but the actual/chance co-occurrence ratio tapered off towards chance level (Fig. 1*H*, Fig. 1-1*C*). Remarkably, the co-occurrence likelihood of bilateral or aHC+pHC HC-SWR remains in an above-chance plateau until the time window in which co-occurrence is measured expands beyond ∼100 ms for bilateral HC or beyond ∼500 ms for a/pHC. This is similar to the window wherein HC-SWR interact with NC-GE, and thus defines a brief period for interaction of multiple HC and NC sites.

Patel et al. (Patel et al., 2013) found in rodents that dorsal SWR often propagate to ventral, but ventral SWR often remain isolated. Accordingly, we tested if the probability of an aHC-SWR given a pHC-SWR was greater than the probability of a pHC-SWR given an aHC-SWR; When tested across all 8 pairs of sites we found this to be a trend, but not significantly so (one-tailed paired t-test, p = 0.149). However, when testing individual sites, we found that significant directionality is present in 7 of 8 patients when both sites generate SWR within the same 200 ms window (Fig. 1*E*, Fig. 1-1*A*). While for 4 of these 8 patients, aHC-SWR preceded pHC-SWR (binomial test, combined N2+N3, p < 2.20 × 10^−16^ for patient 4, p = 1.72 × 10^−10^ for patient 5, p < 2.20 × 10^−16^ for patients 9 and 19), the opposite was found for 3 patients (p < 2.20 × 10^−16^ for patient 6, p = 5.79 × 10^−11^ for patient 11, p = 2.80 × 10^−8^ for patient 13), and 1 showed no significant preference either way (patient 3, p = 0.2082) (Fig. 1*E*, Fig. 1-1*A*). Thus, SWR in most pairs of HC sites show significant order effects, but these are not consistently in the anterior-to-posterior or posterior-to-anterior direction.

The dominant and non-dominant hippocampal formations of humans are thought to be specialized for verbal and visuospatial memory, respectively (Glosser et al., 1998). This might predict distinct memory traces and thus low bilateral co-occurrence of SWR. We tested this prediction in patients (n = 4) with bilateral (left and right) HC contacts, both placed in either anterior or posterior HC. In these patients, co-occurrence between HC in different hemispheres was 2.4±1.2% in N2, and 2.8±1.2% in N3. While this was less than that usually observed between two contacts within the same HC, it was still significantly greater than chance-derived overlap percentages (1.1±0.64% in N2, p = 0.0317; 1.4±0.77% in N3, p = 0.0105, one-tailed paired t-test) (Fig. 1-1*B*).

Thus, in humans, only a small proportion of SWR co-occur in both anterior and posterior hippocampal regions, and an even smaller proportion in both left and right hippocampi. This general independence of HC-SWR from different longitudinal or bilateral origins indicates that any particular HC-SWR only engages a small proportion of the HC.

### HC-SWR are significantly associated with all categories of neocortical graphoelements (NC-GE), including theta bursts (TB), downstates (DS), upstates (US), and sleep spindles (SS)

In order to examine the relationship between HC-SWR and NC-GE, different categories of GE (TB, DS, US, and SS) were identified in simultaneous recordings from NC in all lobes. NC recording channels were differential derivations between adjacent contacts spanning the cortical ribbon, and were thus assured of being locally-generated. Detection algorithms have all been extensively validated in our previous studies (|Mak-McCully et al., 2015, 2017; Gonzalez et al., 2018). Down-to-upstate transitions (D2U) were also identified. However, the results for the D2U event were largely redundant with those for DS and US, and thus are only included in Extended Data Fig. 3-2.

In order to display the regularities in our data, we constructed histograms of the occurrence times of each GE (TB onset, SS onset, DS peak, and US peak) between each pair of (non-epileptogenic) HC (n=32) and NC (n=366) channels (n=20 patients). Separate analyses were carried out for stages N2 vs. N3, for SWR recorded in aHC vs. pHC, and for HC-NC pairs in the same vs. opposite hemispheres. Detailed breakdown of histogram contents by GE types and SWR sources (N2 vs. N3, aHC vs. pHC, ipsilateral vs. contralateral) can be found in Extended Data Figure 2-1.

Of the 598 HC-NC pairs, 551 (92%) of all pairs had at least one histogram showing a significant relationship between the HC-SWR and a NC-GE (p<.0.05 post FDR-correction, randomization test, see section “Experimental design and statistical analysis” in Methods). The proportion was slightly greater when the HC and NC sites were in the same hemisphere (432 (94%) of the ipsilateral pairs and 119 (85%) of the contralateral pairs). Of the entire set of 4784 histograms, 1722 (36%) showed a significant association between the HC-SWR and NC-GE. The significant histograms (examples are shown in Fig. 2) include all types of cortical GE from channels across all cortical lobes. The peak of the histogram represented a mean increase of 71%, or median increase of 49% above baseline (measured as the mean height of non-significant bins). We summarized our HC-NC association data further by constructing histograms of peak latencies from the histograms described above. Specifically, we determined the latency of peak occurrence rate relative to HC-SWR time for each significant HC-SWR/NC-GE histogram, and plotted these as a histogram for each GE, separately for N2 vs. N3 and aHC vs. pHC (Fig. 3, Extended Data Fig. 3-1). Results were also summarized by plotting each cortical channel after morphing to a standardized brain (Fig. 3, Extended Data Fig. 3-1).

In the summary histograms, significant temporal relationships were found between HC-SWR and each of the NC-GE types. As shown in Table 2, we tested for the presence of a significantly non-random distribution of NC-GE peak times with respect to HC-SWR (column 4), and found significant results for both N2 and N3, and both aHC and pHC, for TB, DS, and US. NC-SS and HC-SWR were significantly related in N2 pHC with a trend in N3 pHC, reflecting both the preferred occurrence of NC-SS during N2, and a tighter coupling between NC-SS and pHC as compared to aHC, which is described in Jiang *et al*. (submitted-2).

### NC-TB typically precede, while NC-US and SS typically follow HC-SWR

We also examined if the peak latency in the summary histograms between HC-SWR and NC-GE differed significantly from zero, using both the mean of the included HC-NC channel pairs (column 5 of Table 2), or a fitted distribution (column 6 of Table 2). Significant effects were found for SS in N2, which followed HC-SWR by 169 ms (mean) or 244 ms (fitted). DS were also significantly later than HC-SWR in the pHC by 157 ms (mean) or 290 ms (fitted). Finally, US were significantly later than HC-SWR in both N2/N3 and a/pHC by 112 ms (mean) or 179 ms (fitted). The same NC-GE (DS, US and SS) each also showed a significantly greater proportion of NC-GE events in the 500 ms after versus before the HC-SWR (columns 7 and 8 of Table 2). In contrast, the proportions of TB were significantly greater before versus after HC-SWR (column 8 of Table 2). However, when these effects were examined individually for different sleep stages and hippocampal origins, only trends were observed for pHC-SWR (column 7 of Table 2), possibly due to the relatively low count of NC sites showing significant NC-TB to HC-SWR coupling (Fig. 3, Fig. 3-1).

Different types of NC-GEs had different typical latencies with respect to HC-SWR; for example, TB tended to precede pHC-SWR, and spindle onsets tended to follow pHC-SWR by ∼300 ms. Overall, the latencies of different NC-GEs with respect to SWRs shown in Fig. 3*A-C* and Table 2 were consistent with the typical sequence we established in our previous studies of cortical NC-GE during NREM sleep (|Mak-McCully et al., 2015, 2017; Gonzalez et al., 2018). In N2, this sequence begins with the NC-TB which increases in strength to terminate as the negative peak of the NC-DS, and the NC-SS then begins on the upslope to the NC-US. An example of a canonical sequence is shown in Fig. 4*A* together with an HC-SWR between the NC-DS and NC-SS. There are, however, nuances alongside this general trend. In particular, NC-DS peaks significantly occurred after pHC-SWR but not after aHC-SWR, and the NC-SS onsets follow HC-SWR in N2 but not in N3 (Table 2). These apparent discrepancies could be addressed by considering that we identified (and computed histograms for) TB and SS events by their onsets, while DS and US events were identified by their peaks (Fig. 3*D*). Thus, in the case of NC-DS, if DS peaks co-occur with aHC-SWR and slightly follow pHC-SWR, then the actual starts of the DS events would likely precede aHC-SWR (as seen in Fig. 4*A*) and more or less co-occur with pHC-SWR. In the case of NC-SS, while some SS onsets may briefly precede HC-SWR, since SS last ∼.5-2 s, most of the SS events would still take place after HC-SWR. Overall, therefore, US and SS clearly follow, TB clearly precede, and DS more or less happen at the same time as HC-SWR.

**Figure 4.**
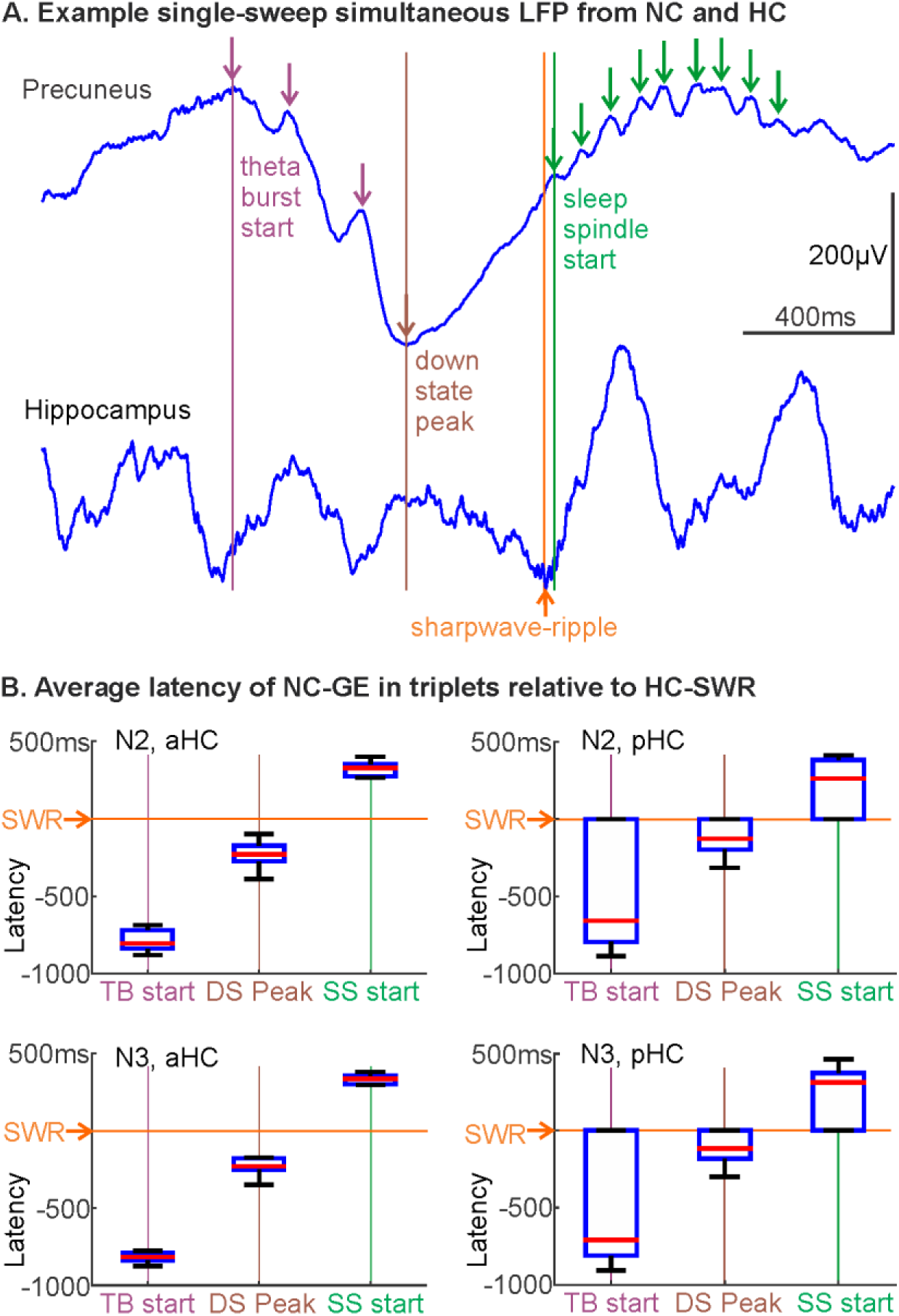
HC-SWR are associated with stereotyped NC-GE sequences. ***A***. Example LFP traces of an aHC-NC channel pair where TB, DS, SWR, and SS occur in sequence. Vertical colored lines indicate automatically detected NC-GE locations (TB start, DS peak, SWR center, and SS start). ***B***. In TB→DS→SS sequences that co-occur with HC-SWR, the latencies between NC-GEs and SWRs are stereotyped, with TB starts almost always preceding HC-SWR and SS starts almost always following.

Although generally consistent with the TB-DS-SS/US canonical sequence, the temporal ordering of NC-GE would be obscured by the fact that they are not necessarily associated with the same SWRs. Thus, we performed an analysis limited to events wherein NC-TB, NC-DS and NC-SS all occurred in proximity (−1500 ms to +1000 ms) to the same HC-SWR. We found that the canonical order (NC-TB→NC-DS→NC-SS) was about twice as common as other orders (p=0.0001, two-tailed paired t-test for in-category vs out-category over all NC channels, combining N2/N3/aHC/pHC), with TB-DS-SS occurrence rate in NREM being 14% higher than the rate of other possible sequences (0.40/min vs. 0.35/min). Furthermore, in the canonical sequences, the latencies of the successive NC-GE and HC-SWR were highly stereotyped (Fig. 4*B*), especially in the events anchored by an aHC-SWR. Since SS starts and US peaks tend to overlap in their latency distributions with regard to HC-SWR, we repeated the canonical order analysis with SS and US combined (so a typical sequence would be TB-DS-SS/US), and found the NC-TB→NC-DS→NC-SS/US to be 25% more common than other orders (p = 0.0339), as expected.

### Multiple NC-DS are more likely than single NC-DS to co-occur with HC-SWR

In previous studies with SEEG, we showed that neocortical downstates (NC-DS) vary in the extent of cortex which they engage, with the NC-DS involving greater cortical extents being more likely to evoke a thalamic DS (|Mak-McCully et al., 2015, 2017). Thus, the broadband spectral power decrease we see in Fig. 1*D* is suggestive of a DS-like state in the HC, i.e. the NC-HC relationship may be similar to the NC-thalamic interactions previously examined. We therefore predicted that HC-SWR may be associated with the convergence of NC-DS from multiple cortical locations. Indeed, we found that multiple (i.e. more than 1, up to 9 in our data) NC-DS are significantly more likely than single DS to co-occur with HC-SWR from N2 pHC (15% increase), N3 aHC (13% increase), and N3 pHC (18% increase) (p = 0.0021, 0.0026, and 0.0029, respectively, one-tail paired t-tests). In the case of N2 aHC, there was a trend for multiple NC-DS to co-occur with HC-SWR more often (5% increase, p = 0.075, one-tail paired t-tests).

To ensure that this increased likelihood of HC-SWR co-occurrence with multiple NC-DS cannot be explained by patient-wise variability alone, we built a linear mixed-effect model (with aHC/pHC and N2/N3 data amalgamated) of the SWR-DS co-occurrence rate, using “single” vs. “multi”-DS categories—i.e. whether a given HC-SWR co-occurs with a single DS or more than one DS—as the fixed effect, while accounting for patient-wise variability as random effect (intercept). As expected, we observed a significant effect of single vs. multi-DS on SWR-DS co-occurrence in this model (p = 0.0013). Thus, the tendency of HC-SWR to co-occur with multiple NC-DS cannot be reduced to sampling error, and may represent a physiological feature possibly underlying synchrony across NC, HC, and thalamus.

### The NC sites whose GE co-occur with SWR in a given HC site are not random, but specific for the HC location

The results presented above demonstrate that for a given HC site, a particular category of GEs will significantly co-occur in some but not other NC sites. Indeed, the striking positive skewness both in general and in specific categories (Fig. 2-1) suggests that, in addition to high variability in co-occurrence rates of NC-GEs with SWRs, a small population of histograms for particular HC-NC-GE-state combinations (examples shown in Fig. 2) may contain a disproportionately large number of events. However, they would not bias the main results (Figs. 3, 3-1) where each of the significant HC-NC channel pairs provide a single count.

We further examined this issue by performing a chi-square test of homogeneity for each HC site with regard to co-occurrence of each NC-GE, across NC sites. We found that most HC sites were coupled to NC-GE with NC site distributions that significantly differ from chance (Fig. 3-3): out of 32 unique HC sites, 21 were coupled to non-homogeneous distributions of NC sites with significant TB-SWR relationships, 21 were coupled to significant SS-SWR NC site distributions, and all 32 were coupled to significant DS and US distributions. NC sites where GE co-occur with HC-SWR from a particular HC location constitute an anatomical network which could have functional significance during consolidation, though limitations of our current study necessitate further support via behavioral assessment in future investigations.

Having determined above that the HC-NC co-occurrences are not randomly distributed across the recorded NC sites, we then examined if the distribution differed between HC sites. This analysis was conducted in patients with multiple HC contacts either in the same hippocampus (aHC and pHC) or bilaterally in both anterior or both posterior HC (Fig. 3-4). Again, we repeated this analysis for each of the NC-GE categories (Fig. 3-5), provided that both HC sites had a significantly non-random pattern of co-occurrence with NS sites. For a given NC-GE, analysis was limited to those NC sites that significantly co-occurred with both HC sites (i.e., both aHC and pHC, or both left and right HC, in a given patient). In total, 8 a/pHC pairs and 4 Left-Right HC pairs were examined. We found a tendency for aHC/pHC pairs to yield significantly different HC-NC coupling distributions, with NC-DS and NC-US showing stronger effects than NC-TB and NC-SS (Fig. 3-4). Similarly, for Left-Right HC pairs, all 4 of the qualified pairs for NC-TB, NC-SS, and NC-US showed significant differences, and all 3 of the qualified pairs did for NC-DS (Fig. 3-4).

We then repeated this procedure, but instead of comparing proportions between HC site pairs, we compared between NC-GE types for each HC site (Fig. 3-5). Each test required that for a given HC site, both NC-GE types needed to have a significantly non-random co-occurrence pattern across the NC sites, as described above (Fig. 3-3). For two of the six possible NC-GE site pairs (TB vs. SS, DS vs. US), we found remarkably few HC sites showing a significant difference. Thus, in a particular patient, if a given HC-NC site-pair is significantly related for SWR-TB, then it will probably also be related for SWR-SS. Similarly, if a HC-NC site-pair is related for SWR-DS, then it will probably also be related for SWR-US. Other pairs of NC-GE’s (TB vs. DS, TB vs. US, and SS vs. US) did not show this correspondence, with another (SS vs. DS) showing a mixed pattern.

In summary, the particular pattern of NC sites whose GE co-occur with SWR from a given HC site is not random, but is specific for the HC location and GE (although SS/TB and DS/US show similar patterns).

### HC-SWR co-occur with NC-GE more often for DS/US than TB/SS, in NREM sleep stage N2 than N3, in posterior than anterior HC, and in frontal than occipital regions

While the summaries in Fig. 3 and Table 2 gave us an overview of NC-GE and HC-SWR relations, we noted that the maps in Fig. 3 revealed apparent regional differences across NC in co-occurrence strength. We also considered the possibility that “hot spots” may exist in NC and contribute to the widespread positive skewness seen in Fig. 2-1. We therefore explored the spatial distribution of HC-NC relationship further by tallying the proportion of significant HC-NC channel pairs across different NC regions, with respect to different NC-GE types and different HC-SWR sources (Table 3). To explore the apparent variability in Table 3, we performed a 4-way ANOVA to compare the main effects of GE type (TB, SS, DS, US), NREM stage (N2 vs. N3), a/pHC origin of HC-SWR, and NC ROIs (coverage listed in Table 3 and Table 3-1), as well as the 2-way and 3-way interaction effects (Table 4). All post-hoc analyses were performed with Tukey’s range test. We checked the normality assumption by conducting the Lilliefors test on the residuals, and the null hypothesis of normality could not be rejected (p = 0.1653).

**Table 4.**
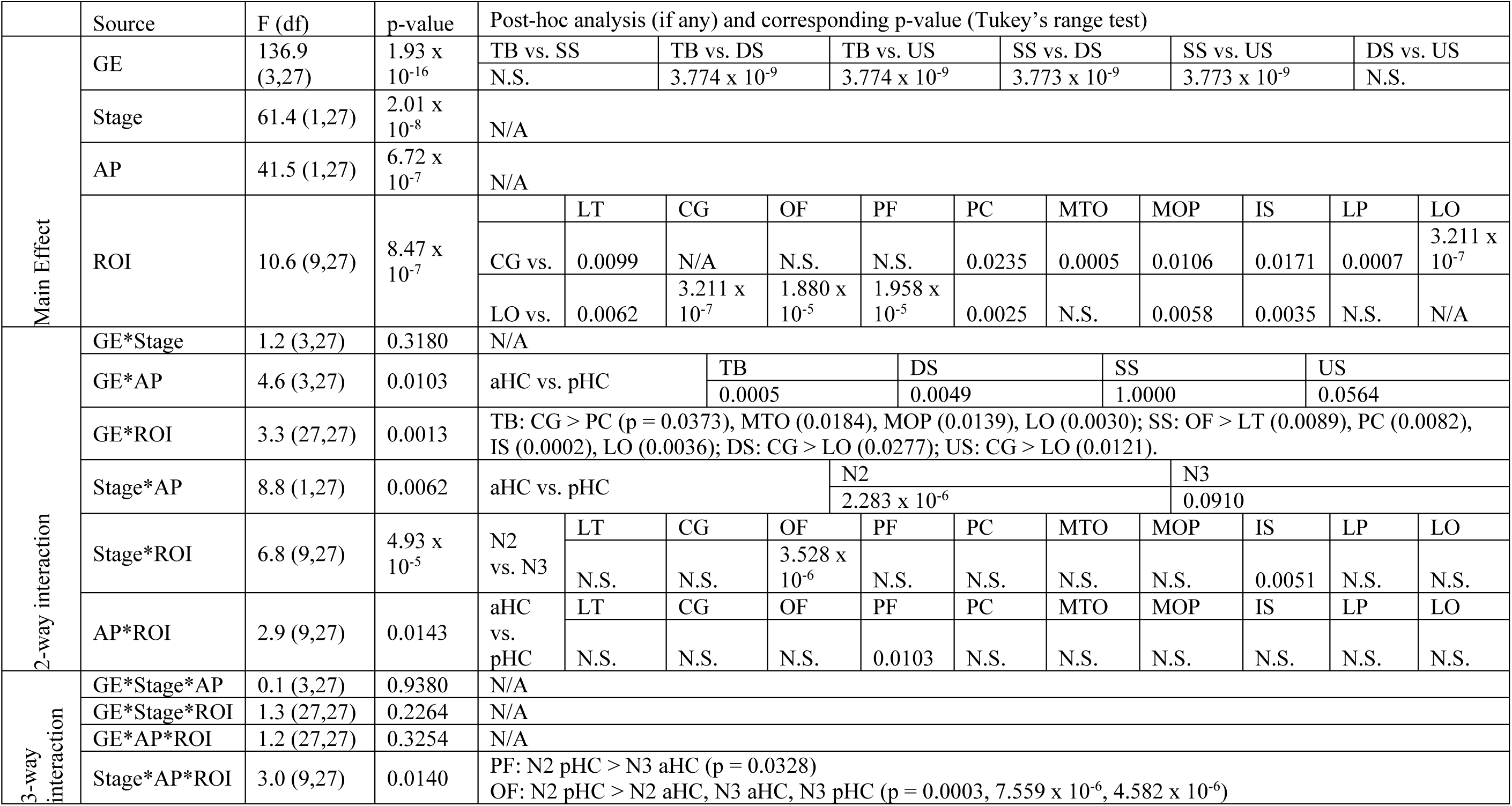
Summary of 4-way ANOVA comparing associations between HC-SWR and NC-GE. For each HC-NC electrode pair, and sleep stage (N2 versus N3), the significance of the NC-GE occurrence times relative to the ripple center was determined (see Fig. 1 and Methods). We then tallied the proportion of significant channel pairs according to their HC location (aHC, pHC), NC location (see Table 3-1 and Fig. 5*A*), and GE type (TB, DS, SS, US). These proportions were then entered into ANOVA. LO: lateral occipital; LT: lateral temporal; CG: cingulate; OF: orbitofrontal; PF: prefrontal; PC: paracentral; MTO: medial temporo-occipital; MOP: medial occipito-parietal; IS: insula; LP: lateral parietal. N.S.: not significant (for post-hoc Tukey’s range tests, p > 0.1). df: degrees of freedom.

All 4 main effects were significant. The GE main effect was due to HC-NC coupling being greater for DS (0.5136) and US (0.5299) than TB (0.1655) and SS (0.1852). The NREM stage main effect was due to higher HC-NC coupling in N2 (0.412) than N3 (0.294) in N3, despite HC-SWR occurring more often in N3 (Table 1). The fact that the durations of N2 were about twice those of N3 in our recordings may have contributed to this unexpected observation. We therefore repeated the 4-way ANOVA for the main effects using data from a subset of 6 patients with similar N2 and N3 durations (patients 1, 6, 9, 10, 19, and 20), and found the NREM stage effect to remain significant (F = 7.77, df = 1, p = 2.006 × 10^−8^); the effect size for the subset ANOVA is smaller compared to the original (partial eta squared, 0.2235 vs. 0.6945), but still large by convention (Cohen, 1969). The aHC/pHC main effect was due to lower coupling with aHC (0.316) than pHC (0.414 for pHC), despite the density of SWR being about twice as high in aHC versus in pHC (Table 1). The NC ROIs main effect was due to an anterior-to-posterior gradient, whereby the more frontal an NC region, the higher the sum tended to be (Fig. 5*A-B*). The exception was cingulate, which had the largest sum overall, and post-hoc analysis across NC ROIs confirmed that, except for orbitofrontal and prefrontal, the mean proportion for cingulate was significantly greater than all other NC ROIs. In contrast, the most posterior ROI (lateral occipital) had the smallest sum overall. Thus, HC-NC coupling occurred in all regions but was generally larger frontally and smaller occipitally.

**Figure 5.**
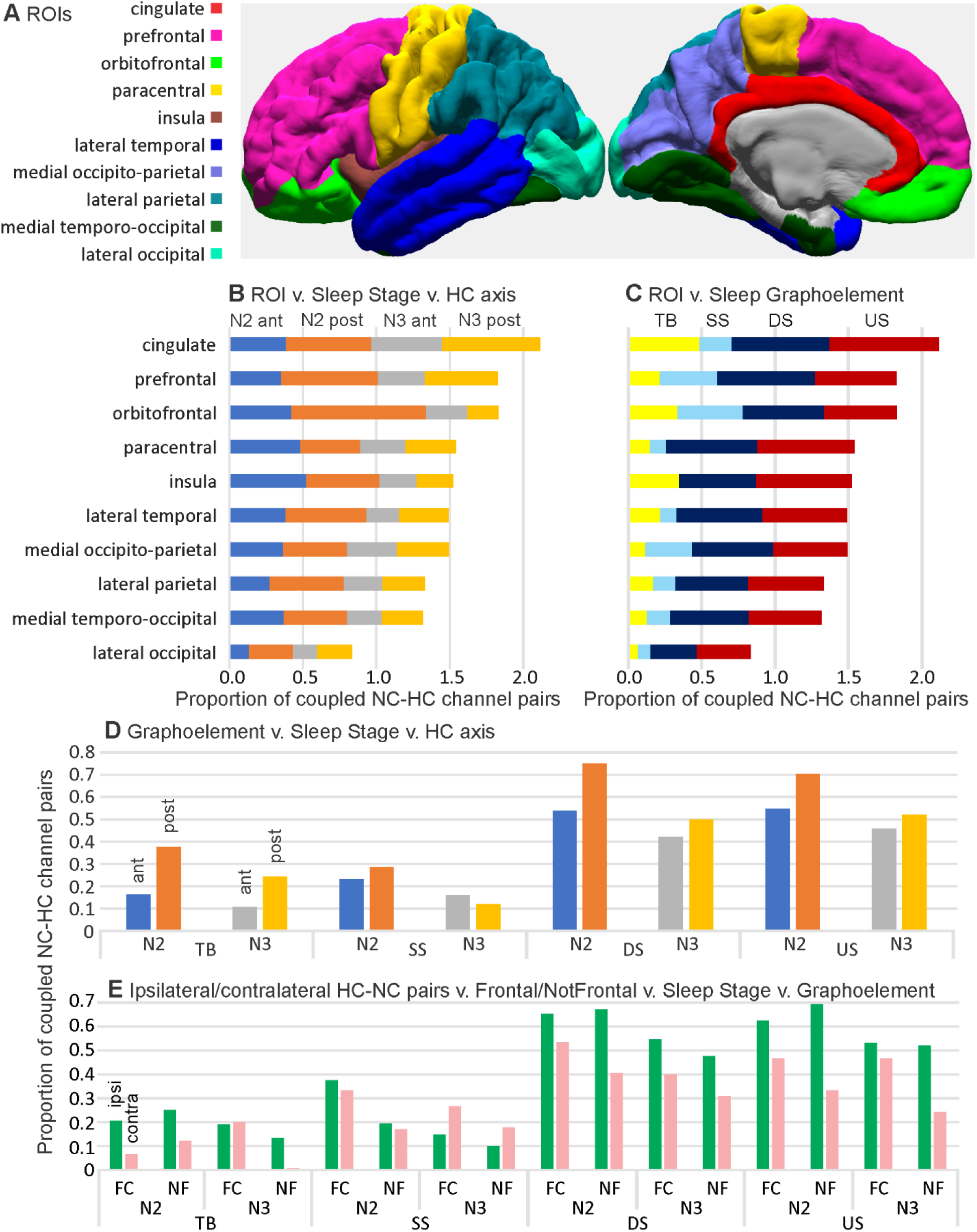
Strength of HC-NC association varies across NC regions (***B***, ***C***, ***E***), NREM stages (***B***, ***D***, ***E***), GE types (***C***, ***D***, ***E***), and anterior vs posterior (***D***) or ipsilateral vs contralateral (***E***) SWR sources. ***A***. Map of NC ROIs. ***B***, ***C***. Mean proportions of significant HC-NC channel pairs in different NC regions. Channels with significant (as evaluated in Fig. 2) HC-NC co-occurrences are more common in fronto-central (FC) than non-frontal (NF) cortex (***B***, ***C***, ***D***), N2 than N3 (***B***, ***D***, ***E***), DS and US than TB and SS (***C***, ***D***, ***E***), with posterior than anterior HC-SWR (***D***), and with ipsilateral than contralateral HC-SWR (***E***). Within these general patterns, interactions can be observed, such as the relatively high co-occurrence of orbitofrontal GE with pHC-SWR in N2 (***B***). Proportions are relative to total channels in each ROI (***B***, ***C***), all ROIs (***D***), or fronto-central or non-frontal ROIs in the ipsilateral vs contralateral hemisphere (***E***). For correspondence of ROIs to FreeSurfer labels, and their amalgamation to fronto-central and non-frontal, see Extended Data Table 3-1.

### NREM stage, GE type, anterior vs posterior HC-SWR origin, and NC ROI, have interacting effects on SWR-GE co-occurrence

Except for GE * NREM stage, all 2-way interaction effects from the ANOVA were significant (Table 4). The GE * aHC/pHC interaction was due to NC-SS only showing pHC-SWR association preference over aHC in N2, but not in N3 (Fig. 5*D*), and thereby did not show significant aHC/pHC difference overall in post-hoc analysis, while other NC-GE types all had pHC preference. The GE * NC-ROI interaction was consistent with our observation that certain NC regions, e.g. insula, cingulate, and orbitofrontal, could be more related to some NC-GE types than others (Table 3, Fig. 5*C*). Post-hoc analysis confirmed this, e.g. for cingulate and lateral occipital ROIs, NC-TB, DS, and US from cingulate were significantly more related to HC than lateral occipital ones were, but the same cannot be said for NC-SS (Table 4). The NREM stage * aHC/pHC interaction was due to a higher coupling advantage for pHC over aHC in N2 than N3, with the exception of NC-SS (Fig. 5*D*). This may be related to the preference of NC-SS for N2 (Andrillon et al., 2011). The NREM stage * NC ROI interaction was due to orbitofrontal and insula having greater proportions of significant HC-NC coupling channels in N2 than N3, while other ROIs showed no preference. Notably, the proportion of orbitofrontal channels with significant SS-SWR relationship was the highest among ROIs (Fig. 5*C*), possibly contributing to orbitofrontal’s N2 preference as described above. The aHC/pHC * NC ROI interaction was due to a preference for pHC by prefrontal sites, and the lack of significant aHC/pHC preference in all other ROIs, emphasizing the widespread NC coupling to SWR in both aHC and pHC (Fig. 3, Fig. 5).

As previously mentioned, the interaction effect of GE * NREM stage was not significant, despite the fact the NC-TB and NC-SS occur more often in N2 (Andrillon et al., 2011; Gonzalez et al., 2018), while NC-DS and NC-US occur more often in N3 (Cash et al., 2009). This observation suggested that the GEs which did occur in the sleep stage where they occurred less often overall might not change in how related they were to the HC-SWR, i.e. the HC-NC relationships (as estimated from GE-SWR correlations) were consistent over time in NREM. Among the 3-way interaction effects, only NREM stage * aHC/pHC * NC ROI was significant, reflecting the prefrontal preference for pHC, and orbitofrontal preference for N2.

### NC-GE co-occurrence with HC-SWR is stronger within hemisphere than between hemispheres

Given the well-known lateralization of human hippocampal function (Glosser et al., 1998), we were interested in examining whether ipsilateral HC-NC channel pairs would show different relationships from contralateral pairs across NC-GE types, NREM stages, or NC ROIs. Due to a sparse representation of contralateral channel pairs for individual ROIs, we calculated proportions for larger regions by combining both HC and NC sites. Anterior and posterior HC channels were combined into a single HC category. The 10 NC ROIs used in previous analysis were collapsed into two: “fronto-central”, which included the previous cingulate, orbitofrontal, prefrontal, and paracentral ROIs; and “non-frontal”, which included the previous lateral temporal, medial temporo-occipital, medial occipito-parietal, insula, lateral occipital, and lateral parietal ROIs. We then computed for each ROI the proportion of NC channels significantly coupled to HC-SWR (Table 5), and performed 4-way ANOVA as previously described, with the ipsilateral/contralateral factor replacing the aHC/pHC factor. All post-hoc analyses were performed with Tukey’s range test. We checked the normality assumption by conducting the Lilliefors test on the residuals, and the null hypothesis of normality could not be rejected (p = 0.0950).

**Table 5.** Proportions of HC-NC channel pairs with significant GE-SWR relationships across NC, separated by ROI, NREM stage, GE type, and ipsilateral/contralateral HC-NC pairings. ipsi: ipsilateral. contra: contralateral.

|  | Proportion of channels with significant HC-SWR co-occurring with NC-GE types |  |  |  |  |  |  |  |  |  |  |  |  |  |  |  |
| --- | --- | --- | --- | --- | --- | --- | --- | --- | --- | --- | --- | --- | --- | --- | --- | --- |
| GE Type | TB |  |  |  | DS |  |  |  | SS |  |  |  | US |  |  |  |
|  | N2 |  | N3 |  | N2 |  | N3 |  | N2 |  | N3 |  | N2 |  | N3 |  |
| ROI \ SWR Source | ipsi | contra | ipsi | contra | ipsi | contra | ipsi | contra | ipsi | contra | ipsi | contra | ipsi | contra | ipsi | contra |
| Fronto-central | 0.206 | 0.067 | 0.191 | 0.200 | 0.652 | 0.533 | 0.546 | 0.400 | 0.376 | 0.333 | 0.149 | 0.267 | 0.624 | 0.467 | 0.532 | 0.467 |
| Non-frontal | 0.251 | 0.122 | 0.134 | 0.008 | 0.671 | 0.407 | 0.476 | 0.309 | 0.195 | 0.171 | 0.101 | 0.179 | 0.694 | 0.333 | 0.521 | 0.244 |

All 4 main effects were significant. As our previous ANOVA already revealed significant main effects for GE type and NREM stage, we focused our attention on the remaining two factors (i.e. ipsilateral/contralateral and combined NC ROI). The main effect of ipsilateral/contralateral channel pair types yielded F_(1,3)_ = 65.058, p = 0.0040. and together with mean proportions of 0.395 for ipsilateral and 0.282 for contralateral (a 40% increase), demonstrated that NC-GE in more channels significantly co-occurred with ipsilateral HC-SWR than contralateral HC-SWR. The main effect of ROIs yielded F_(1,3)_ = 28.267, p = 0.0130, indicating that the more frontal/cingulate parts of NC were significantly more likely to show significant GE-SWR correlations than the posterior regions (mean proportion of 0.376 for fronto-central versus 0.301 for non-frontal).

Of the 2-way interaction effects, only laterality * GE and laterality * NC ROI were significant. No 3-way interactions were significant. Post-hoc analysis of the significant laterality * GE interaction (F_(3,3)_ = 14.98, p = 0.0261) revealed a stronger laterality effect for NC-US (p = 0.0287), and a trend for DS (p = 0.0514). Post-hoc analysis of the significant laterality * ROI interaction (F_(1,3)_ = 10.450, p = 0.0481) indicated a stronger laterality effect for the non-frontal ROI (p = 0.0124).

These statistical results are reflected in the proportions of ipsilateral versus contralateral HC-NC channel pairs with significant HC-SWR coupling for each NC-GE type (Fig. 5*E*). Generally, for DS and US, the proportion of significant HC-NC channel-pairs was high ipsilaterally (.58 and .60), lower but still substantial for contralateral (.37 and .31), with consequent contra/ipsi ratios of .64 and .51. TB and SS were both low for ipsilateral HC-NC channel-pairs (.19 and .18), but TB dropped contralaterally (.07) while SS remained constant (.19), resulting in contra/ipsi ratios of .37 and 1.0. Thus, there was a remarkable and specific lack of ipsi/contra-lateral difference in the proportion of NC sites whose SS were correlated with HC-SWR. Overall, the proportion of ipsilateral channels whose NC-GE significantly co-occur with HC-SWR is 40% greater than contralateral, and this effect is greater for NC-US and non-frontal locations.

## Discussion

Overall, our study demonstrates that human HC-SWR are significantly associated with all categories of NC-GE in all recorded NC areas, but for any given HC site only includes a minority of the cortex. The intensity and topography of HC-NC coordination differ across GE category, NREM sleep stage (N2 versus N3), HC location (anterior versus posterior), and NC area (frontal versus posterior, ipsilateral versus contralateral). In addition, HC-NC interactions are temporally-organized, with HC-SWR tending to follow TB onsets (Fig. 6*A*), occur during DS (Fig. 6*A-B*), and precede SS/US (Fig. 6*C*); while DS peaks and SWR have overlapping latencies, the DS initiation typically precedes SWR onsets (Fig. 3). We discuss here how these spatio-temporal characteristics of HC-SWR interaction with NC-GE may assist memory consolidation by coordinating information transfer/processing. Caveats in these interpretations include: recordings may include pathological activity despite strenuous efforts to exclude it; cortical and hippocampal sampling is extensive but nonetheless sparse in each individual; and a causal role of HC-SWR in replay has not been demonstrated in humans (but see Zhang et al., 2018). Specifically, in our study, we did not identify the hippocampal firing patterns which replayed an event from the previous day, nor demonstrate that the consolidation for the replayed event was enhanced if the HC-SWR co-occurred with NC-GE.

**Figure 6.**
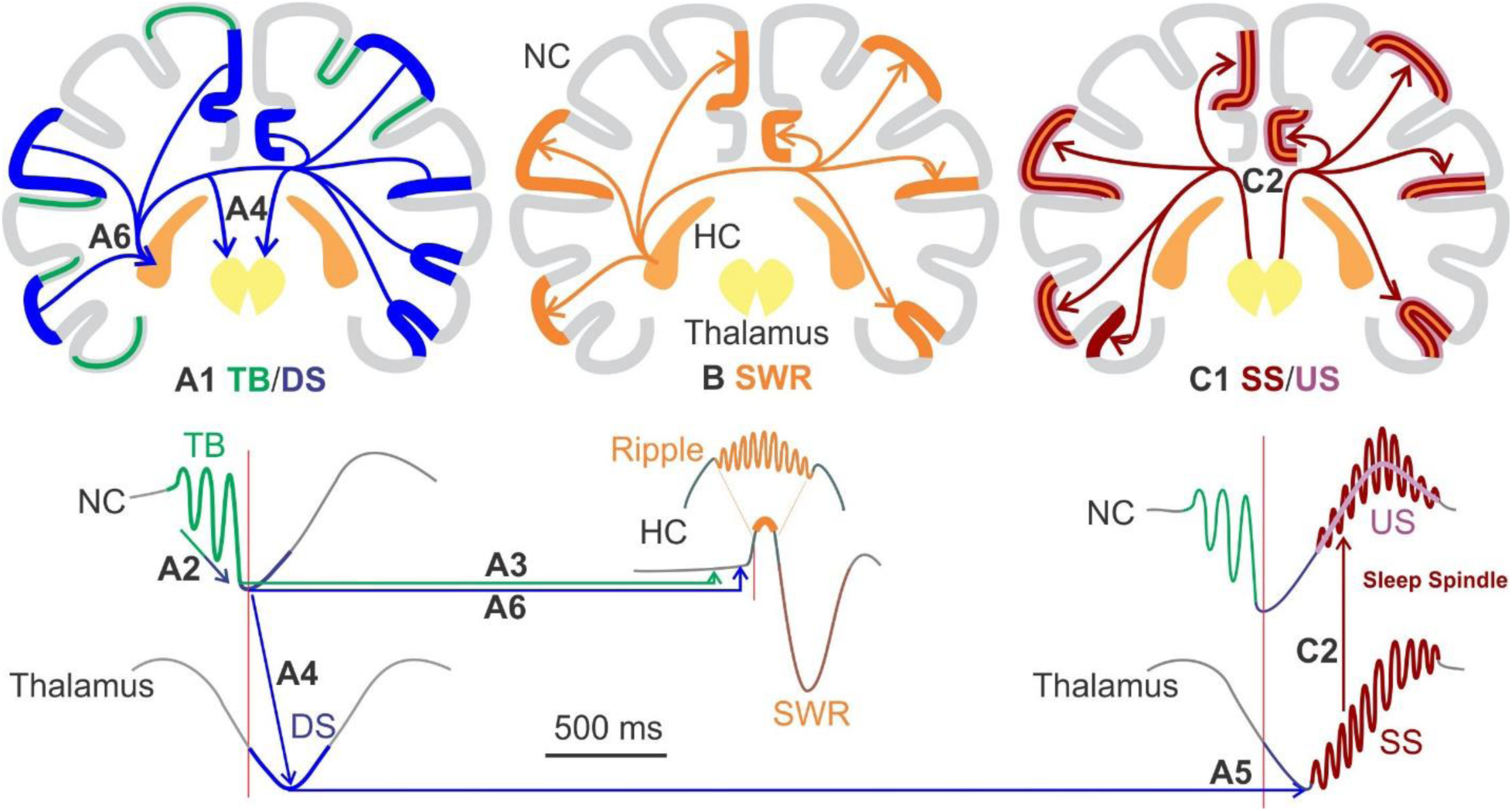
Proposed sequence and interactions of HC-SWR with GE in NC and thalamus. ***A1***. NC-TB appear to arise spontaneously in widespread cortical areas, and may trigger NC-DS (***A2***). TB modulate cortical firing before HC-SWR, which would reflect earlier learned networks, and may be projected to the HC prior to the SWR (***A3***). Thalamic hyperpolarization from converging NC-DS (***A4***), would enable h and T currents, thus triggering the Thalamic-SS (***A5***). NC-DS converging on the HC may contribute to SWR occurrence (***A6***). ***B***. The HC-SWR follows, and if humans are similar to other mammals, marks the wide NC distribution of recent event replay. ***C1***. HC-SWR output arrives at the NC DS→US transition, when the thalamic SS is driving the NC-SS (***C2***). Thus, our results suggest how GE may coordinate, sequence and modulate information transfer between NC and HC to enable memory consolidation. While consistent with current and previous observations (Mak-McCully et al., 2017; Gonzalez et al., 2018), these data are correlational and require mechanistic studies in humans and animals for confirmation.

### Spatial organization of HC-SWR interactions with NC-GE

SWR in both aHC and pHC co-occur with NC-GE of all kinds in all cortical areas (Fig. 5), but with quantitative differences between different GE, areas, hemispheres, and NREM sleep stages. Superimposed on these overall tendencies, the SWR from each HC site co-occur selectively and non-randomly with GE generated in a particular pattern of NC sites (Fig. 6*B*). The modularity of HC-NC co-occurrence networks is further reflected in the very low co-occurrence of HC-SWR in different simultaneously recorded HC contacts that we found, as well as the previously described focality of most cortical SS (Piantoni et al., 2017) and K-complexes (Mak-McCully et al., 2015). The lack of SWR co-occurrence between HC in different hemispheres may be expected given the lack of contralateral responses when the HC is stimulated electrically (Catenoix et al., 2011) However, electrical stimulation of the pHC evokes strong responses in aHC, and *vice versa* (Enatsu et al., 2015), but SWR seldom co-occurred in aHC and pHC. Together, this diversity in HC-NC functional relations argues against models which treat SWR and GE as monolithic global brain states, and for models which posit dynamic engagement of specific HC-NC domains. The functional significance of such modules remains to be examined, but they may allow parallel material-specific processing.

Similar HC-NC networks were engaged by TB and SS, with different, larger networks by DS and US. Frontal-GE co-occurred with HC-SWR ∼25% more than average, whereas the lateral occipital GE co-occurred ∼45% less. The frontal lobe is crucial to declarative memory, especially in forming recollection cues, and integrating context with content (Fletcher and Henson, 2001; Badre et al., 2010). Consistent with our observations, medial temporal efferents in primates target association and prefrontal areas more strongly than early sensory cortex (Van Hoesen, 1995; Suzuki and Amaral, 2004). Similarly, the proportion of ipsilateral channels whose NC-GE significantly co-occur with HC-SWR is 40% greater than contralateral, consistent with the lateralization of human hippocampal function (Glosser et al, 1998).

The proportion of significantly related SWR-GE pairs was 37% higher for SWR occurring in pHC than aHC. In rodents, dorsal HC is essential for spatial memory, whereas ventral contributes to contextual emotional associations (Strange et al., 2014). Correspondences in their distinct external connections imply that the rodent dorsal-ventral distinction corresponds to the posterior-anterior in primates(Strange et al., 2014; Ding and Van Hoesen, 2015). The specific pHC-SWR coupling with NC-GE in cingulate and parietal cortices may reflect this anatomical connectivity. Furthermore, our finding of a higher proportion of pHC coupling with NC-GE may reflect the more direct relation to memory of the homologous rodent dorsal HC.

The proportion of HC-NC channel pairs whose HC-SWR/NC-GE significantly co-occurred was 44% higher in N2 than N3, despite the lower HC-SWR rate in N2 (Table 1). Indeed, the relative isolation of TB-DS-SS/US complexes in N2 (Mak-McCully et al., 2017; Gonzalez et al., 2018) may have rendered their association with HC-SWR easier to parse. In any case, this finding implies that co-occurring HC-SWR with NC-GE are more common later in the night, when N2 predominates. Furthermore, in each ∼90 min sleep cycle, N2 precedes N3, suggesting that the HC-NC engagement may become more selective over the course of each cycle, consistent with previous findings indicating a preference for local active potentiation in N2 and for global homeostatic regulation in N3 (Genzel et al., 2014).

The proportion of HC-NC pairs that were significantly related was three times higher for NC-DS or NC-US than NC-TB or NC-SS. This might suggest that the HC-NC pairs related during TB and SS exchange more specific information whereas the HC-NC pairs related during NC-DS and NC-US reflect more contextual information, and/or provide an overall oscillatory context supporting large-scale timing and synchronization. This interpretation is consistent with the higher density of NC-DS and NC-US than NC-TB and NC-SS (Mak-McCully et al., 2017; Gonzalez et al., 2018) and the greater long-range coherence of their constituent frequencies (Halgren et al., 2018).

Previously we found that NC-US sequences occur during waking in particular sequences across the cortex, and tend to occur more frequently in sleep following the waking period where they were identified, compared to preceding nights (Jiang et al., 2017). These “replaying” events were associated with SWR-like events in the HC of 3 patients. In this study, we did not compare spatiotemporal US sequences from waking to those in sleep. However, we did find that HC-SWRs tend to be followed by multiple NC-US. Further work is needed to test if these spatio-temporal patterns of cortical activation are related to memory representations.

### Temporal organization of HC-SWR interactions with NC-GE

While NC-TB, DS, and SS/US can occur separately from each other, they often occur in that order (Mak-McCully et al., 2017; Gonzalez et al., 2018) (Figs. 3, 6). Typically, the sequence begins with an NC-TB whose oscillations may grow into an NC-DS. Thalamic-DS follow NC-DS, especially when multiple NC-DS are present. Thalamic-SS onset is strongly coupled to the thalamic-DS peak. Thus, when the thalamic-SS is projected back to the NC, it arrives on the upslope from the NC-DS and continues through the NC-US.

We found that within this sequence, NC-TB tend to precede HC-SWR, and thus are positioned to help modulate cortical activities patterns that feed into the HC to possibly cue the event which is replayed by firing during the subsequent SWR in rodents (Fig. 6*A-B*). Several studies have found that waking NC-TB occur during cognitive processing, phase coupled with word memory recall (Halgren et al., 2015; Schreiner et al., 2018) and with neuronal firing, as indexed by high gamma (Canolty et al., 2006; Sato et al., 2014; Alekseichuk et al., 2016). NC-TB when asleep have a similar laminar profile and phase-coupling with high gamma (Gonzalez et al., 2018; Halgren et al., 2018). Unlike NC-SS, NC-TB appear to arise spontaneously in the cortex rather than being driven by the thalamus (Gonzalez et al., 2018). The final inhibitory peak of the NC-TB coincides with the local NC-DS, consistent with NC-DS sometimes being triggered by TB (Gonzalez et al., 2018).

NC-DS peak occurrences following NC-TB would tend to decrease the excitatory NC input to HC. This agrees with our observation that HC-SWR are preceded by locally-decreased broadband amplitude (Fig. 1*D*). Inhibitory interneurons in CA1 play a prominent role in the genesis of HC-SWR (Buzsáki, 2015), and in humans begin to fire prior to the HC-SWR (Le Van Quyen et al., 2008). The global decrease in membrane fluctuations prior to the HC-SWR may help create a more stable background upon which the exceptional synchrony underlying SWR can emerge.

The possibility that NC-DS may promote HC-SWR, together with their role in triggering thalamic-DS/SS, suggests that NC-DS may be crucial in synchronizing activity across different structures (Fig. 6*A*). The tight coupling of thalamic-SS onset to the most hyperpolarized peak of the thalamic-DS suggests that the h and T currents, necessary for spindle generation (McCormick and Bal, 1997), are not chronically available during NREM, but require additional hyperpolarization consequent to decreased excitatory cortical input during NC-DS. This interpretation is supported by the observations that NC-DS consistently precede thalamic-DS, and the probability of thalamic-DS is higher following quasi-synchronous NC-DS in multiple locations (Mak-McCully et al., 2017). Similarly, we found here that HC-SWR were more likely when NC-DS occurred in multiple sites quasi-synchronously, than when they occurred in a single site. The implication that converging NC-DS may trigger DS in the HC and thalamus, and consequently synchronized HC-SWR and thalamic-SS, agrees with the finding that HC-SWR-triggered BOLD in monkeys shows thalamic suppression (Logothetis et al., 2012).

We found that widespread NC-SS/US consistently follow HC-SWR (Fig. 6*C*). During NC-US, cortical cells fire at levels comparable to waking (Steriade, 2001). NC-SS tend to occur on the US, with elevated NC-SS density being recently shown to correlate with targeted memory reactivation in humans (Cairney et al., 2018). There is also a phasic additional increase in firing at SS peaks (Gonzalez et al., 2018), which together with a strong apical dendritic Ca^+2^ influx (Seibt et al., 2017; Niethard et al., 2018) would create the conditions for Spike Timing Dependent Plasticity (Froemke and Dan, 2002). Thus, if, as in rodents, human HC-SWR index the replay by HC pyramidal cells of firing sequences from the preceding waking, then our results imply that the information encoded in these firing patterns arrives in an NC prepared to construct firing patterns and consolidate the consequent networks by virtue of the coincident NC-SS and NC-US. Overall, we found that HC-SWR co-occur with NC-GE in multiple dynamic HC-NC networks.

## Acknowledgements

This work was supported by the U.S. Office of Naval Research’s Multidisciplinary University Research Initiatives Program (N00014-16-1-2829), the National Institute of Mental Health (RF1 MH117155 and R01 MH111437), and the National Institute of Biomedical Imaging and Bioengineering (R01 EB009282). The authors would like to thank the following for their support: John Gale, Qianqian Deng, Charles Dickey, Darlene Evardone, Zach Fitzgerald, Chris Gonzalez, Don Hagler, Milan Halgren, Erik Kaestner, Rachel Mak-McCully, Adam Niese, Burke Rosen, and T. G. Venti.

## Supplementary Material

**Figure 1-1.**
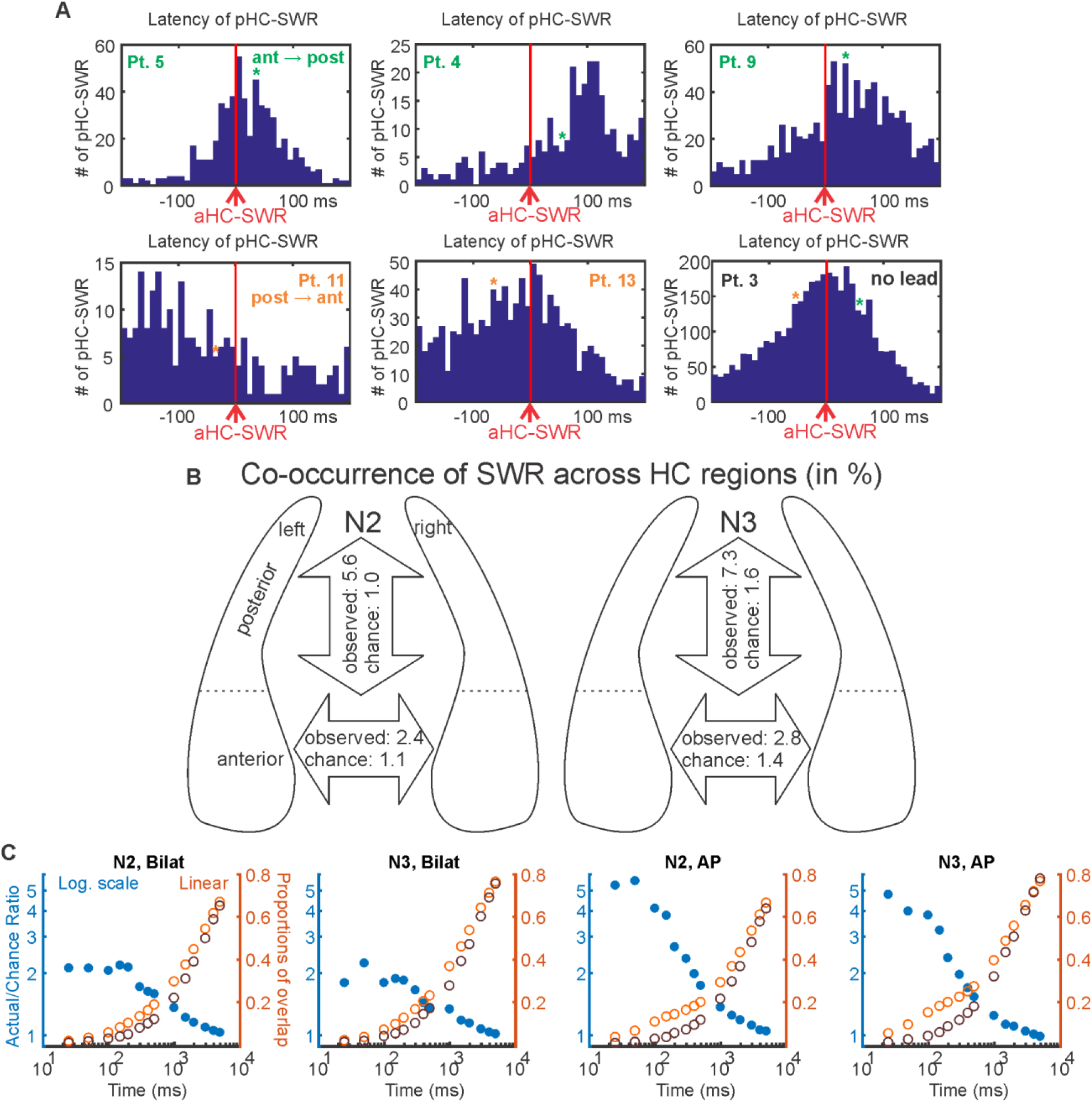
Co-occurrence of SWR from HC site pairs. ***A***. In some patients, aHC SWR tend to precede pHC SWR, while the opposite was found in others. Each panel contains peri-stimulus time histograms over ±200 ms, with aHC SWR as triggers (time 0, red vertical lines). In four patients (marked green), aHC SWR significantly preceded pHC SWR (p < 2.2 × 10-16 for Pt. 4, Pt. 9, and Pt. 19; p = 1.72 × 10-10 for Pt. 5); in three patients (marked orange), pHC SWR significantly preceded aHC-SWR (p < 2.2 × 10-16 for Pt. 6; p = 5.79 × 10-11 for Pt. 11, 2.80 × 10-8 for Pt. 13); in one patient, no significant precedence was observed either way (p = 0.2082) (two-tailed binomial tests for counts within ±200 ms). Pt.: patient. The average occurrence rate of SWRs over NREM (∼0.002 per 10 ms bin). Green/Orange stars mark the time bin closest to when an aHC/pHC SWR would reach pHC/aHC, respectively, given the previously estimated intrahippocampal SWR propagation speed of 0.35 m/s (Patel et al., 2013). ***B***. Sketch of SWR co-occurrence (with co-occurrence criterion being ± 50 ms) across aHC and pHC, as well as across bilateral HCs. The occurrence rates are in numbers per minute. ***C***. For both bilateral HC site pairs and aHC/pHC site pairs. SWR co-occurrence likelihood approaches chance as time window for co-occurrence expands. Blue filled circles mark (wrt. left y-axis) the ratios of actual over chance overlap proportions, and orange/brown unfilled circles mark (wrt. right y-axis) the corresponding actual (orange) and chance (brown) proportions. All axes for each panel are logarithmic. Bilat.: bilateral. AP: aHC/pHC.

**Figure 3-1.**
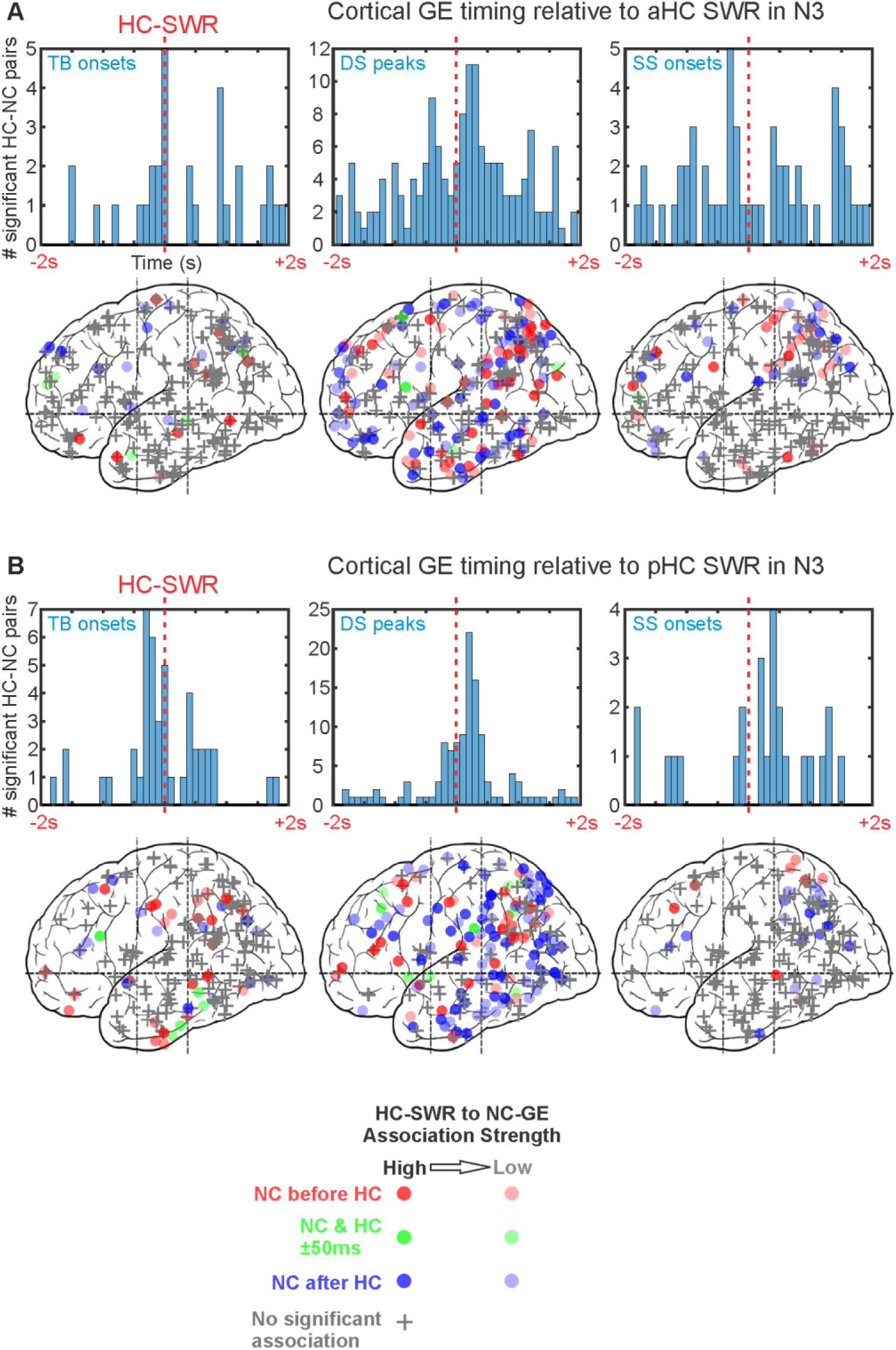
GEs across NC tend to occur with consistent latencies from HC-SWR in stage N3. ***A-B***. Top: Histograms of peak latencies for HC-NC channel pairs with significant temporal correlations between HC-SWR and NC-GE in N3. Each count is the peak latency of a particular HC-NC channel pair. Bottom: maps of peak latency between SWR and GE for cortical channels. Circles indicate where GE-SWR relationships are significant, while plus signs represent non-significant channels. The intensity of each circle corresponds to the strength of the coupling, and the color indicates latency: red for NC-GE before HC-SWR, blue for NC-GE after HC-SWR, and green for NC-GE co-occurring with HC-SWR (i.e. within 50 ms of each other). ***A***. aHC-SWR in N3. ***B***. pHC-SWR in N3.

**Figure 3-2.**
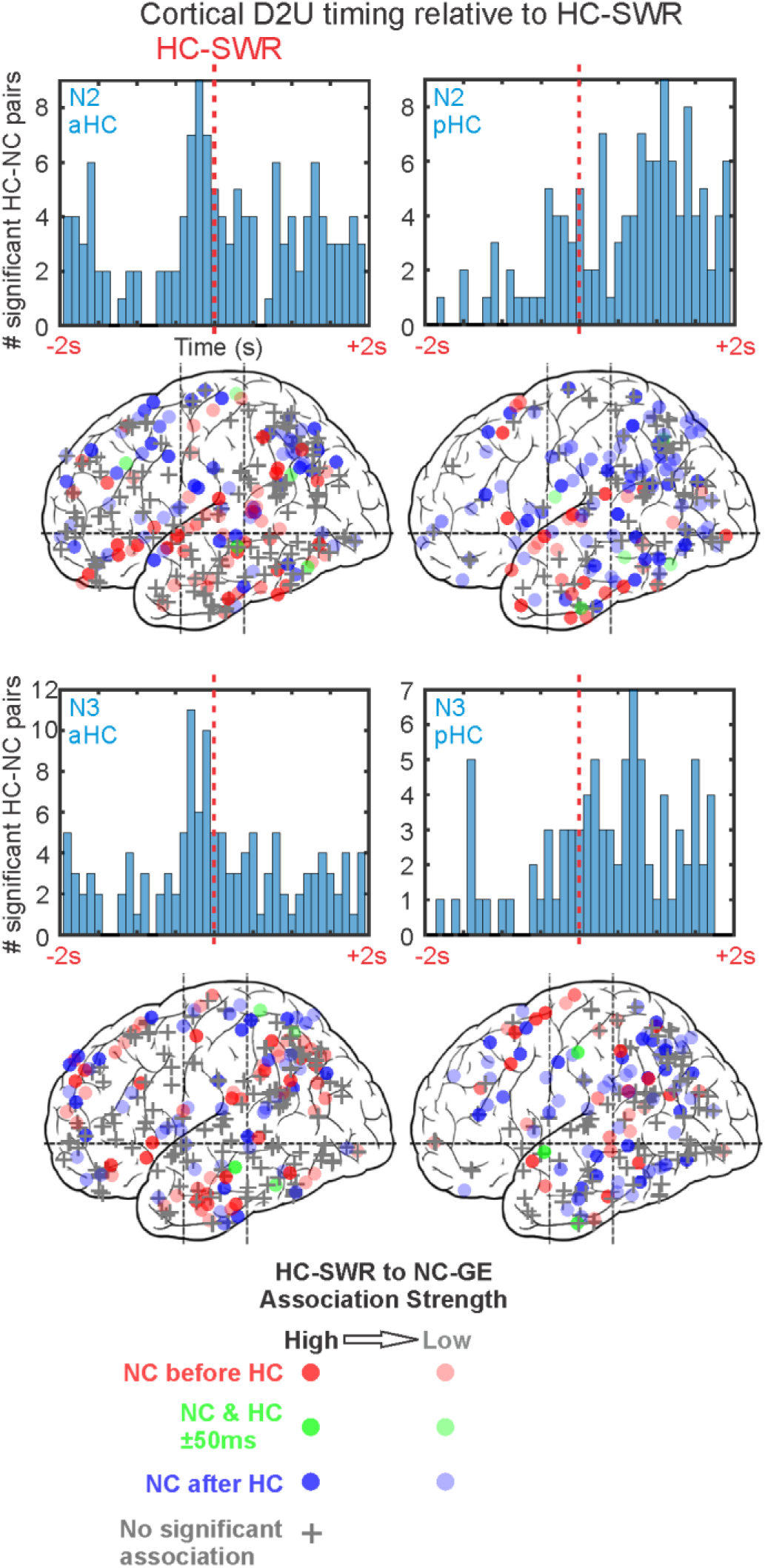
D2U across NC sites co-occur with HC-SWR, but with variable latencies. Top: Histograms of peak latencies for HC-NC channel pairs with significant temporal correlations between HC-SWR and NC-D2U in N3. Each count is the peak latency of a particular HC-NC channel pair. Bottom: maps of peak latency between HC-SWR and NC-D2U for cortical channels. Circles indicate where D2U-SWR relationships are significant, while plus signs represent non-significant channels. The intensity of each circle corresponds to the strength of the coupling, and the color indicates latency: red for NC-D2U before HC-SWR, blue for NC-D2U after HC-SWR, and green for NC-D2U co-occurring with HC-SWR (i.e. within 50 ms of each other).

**Figure 2-1.**
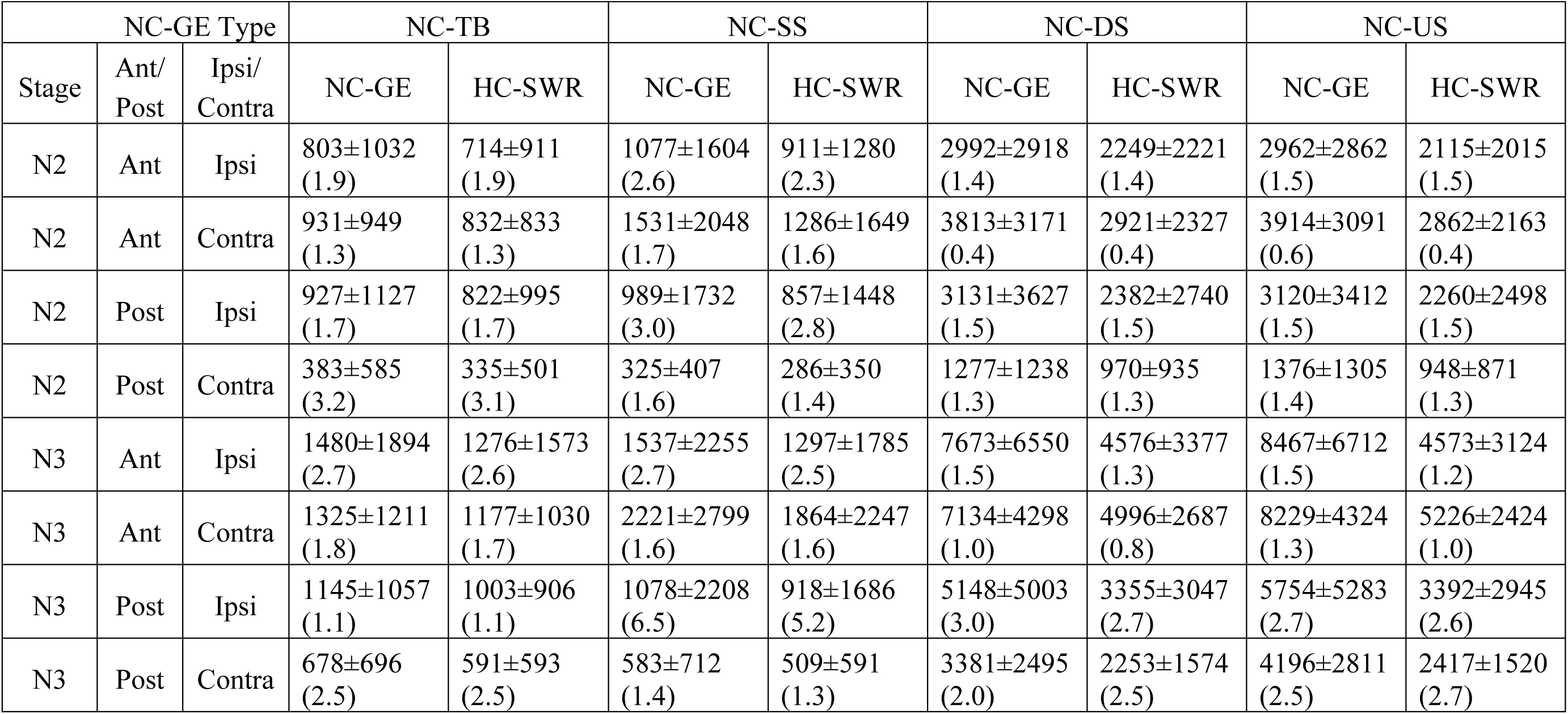
Summary of number of events in NC-GE to HC-SWR histograms. Each cell is formatted as follows: Mean±Standard Deviation (Skewness). Ant: anterior. Post: posterior. Ipsi: ipsilateral. Contra: contralateral.

**Figure 3-3.**
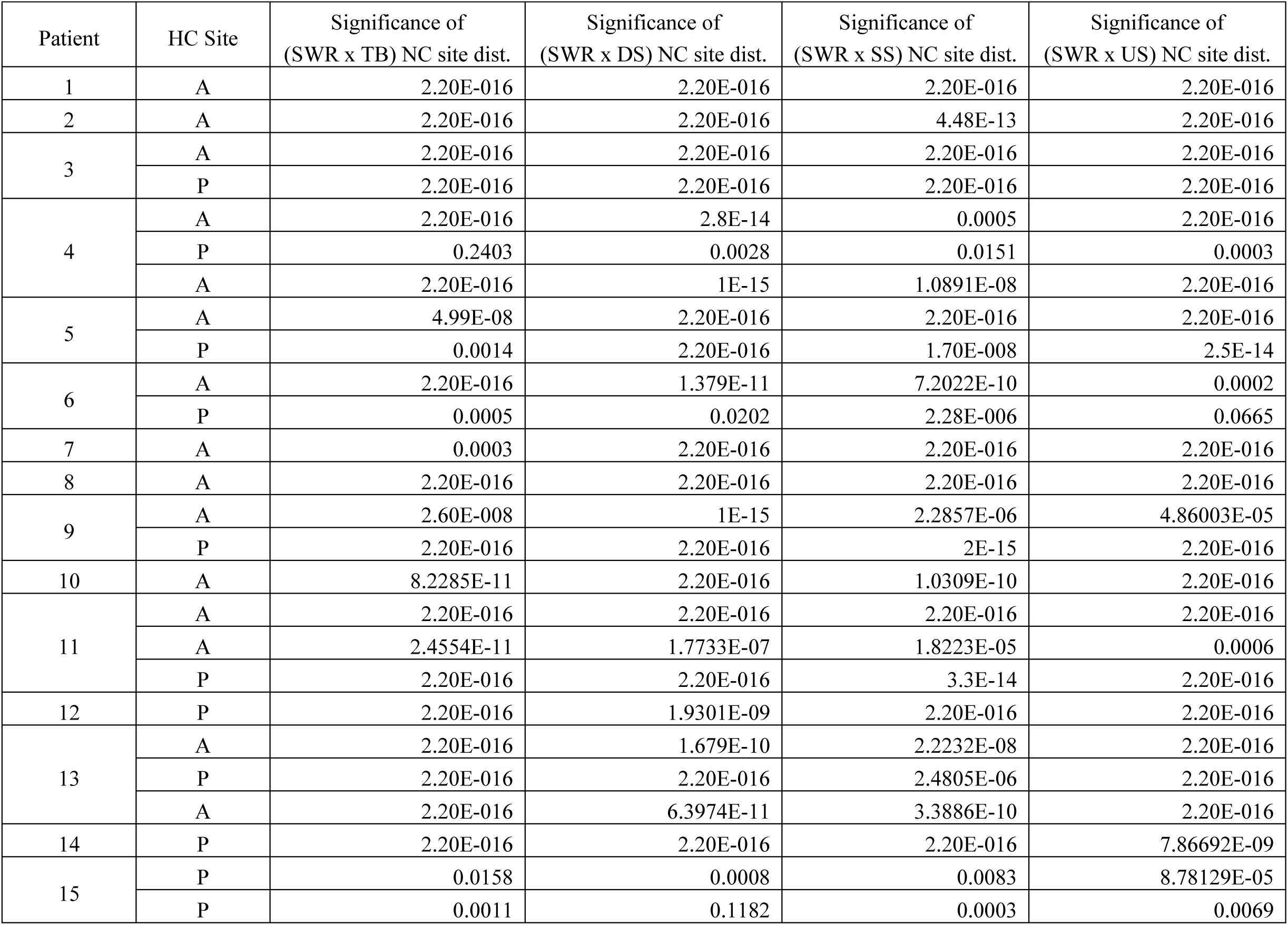
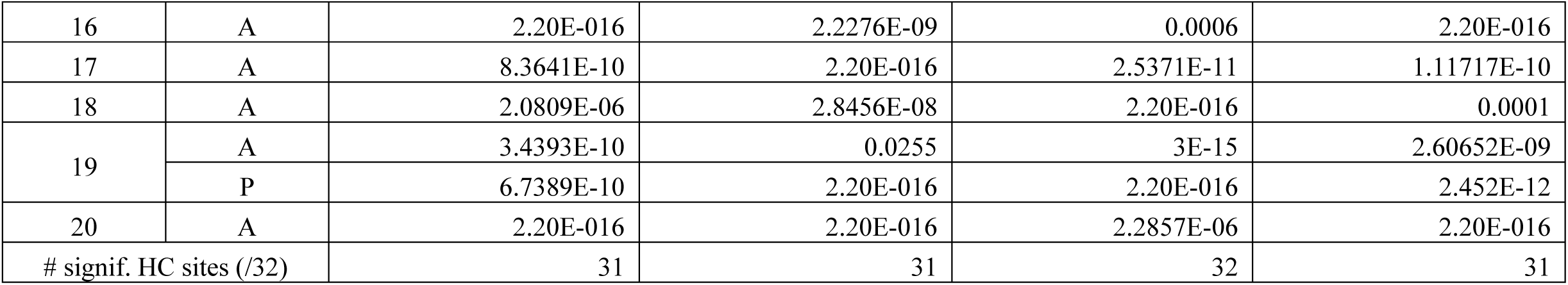
NC-GE synchronized with HC-SWR are not randomly distributed across sites. P-values for chi-square tests of homogeneity (α = 0.05 post FDR-correction applied over each column), conducted for each HC site with regard to each NC-GE’s coupling to HC-SWR in NREM. A: anterior. P: posterior. Cells marked 2.20E-016 are placeholders for p < 2.2 × 10^−16^. # signif.: number of HC sites with a non-homogeneous anatomical distribution of significantly coupled NC sites. Dist.: anatomical distribution (of NC sites).

**Figure 3-4.**
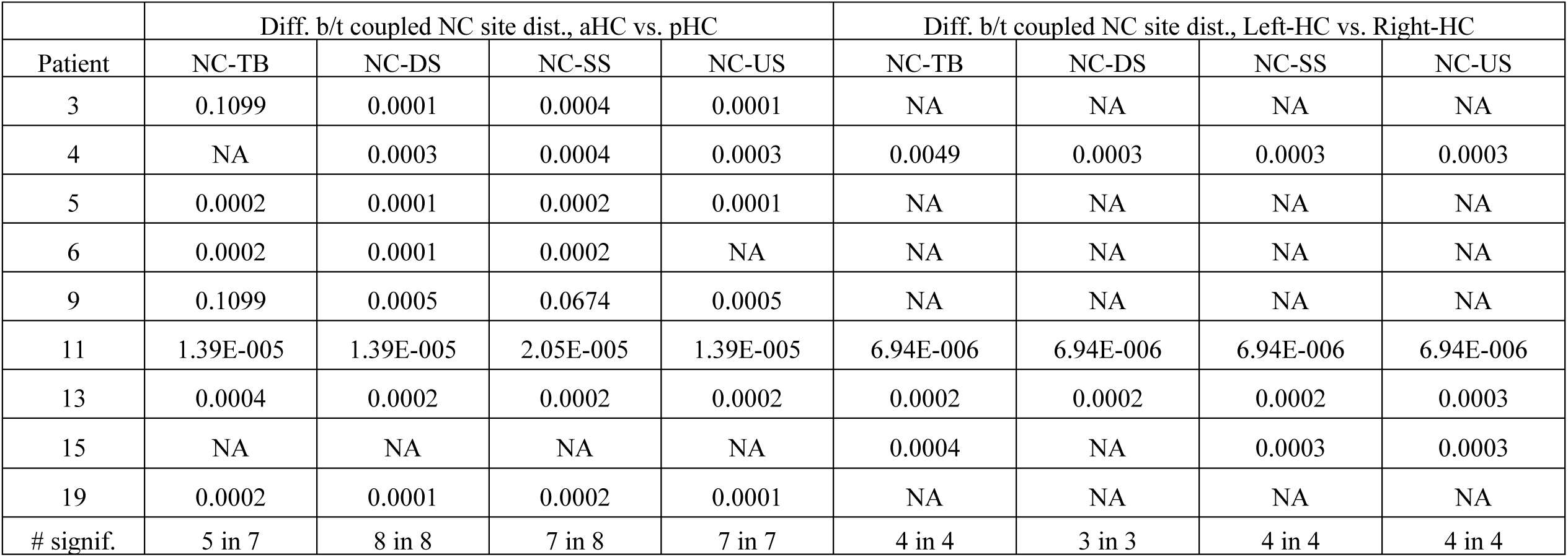
Coupled NC sites differ between simultaneously recorded HC sites. Proportions of NC-GE from significantly coupled NC sites that overlapped with aHC-SWR differ from the proportions for pHC-SWR, and proportions of NC-GE that overlapped with Left-HC SWR differ from the proportions for Right-HC SWR. Wilcoxon signed rank tests were performed on the data obtained for Figure 3-3. Each patient’s test (whose p-value was tabulated above) required that both members of a HC site pair must have significant chi-square test result from Figure 3-3; otherwise, the corresponding cell would be filled with NA. # signif.: number of significant tests (in total number of qualified patients).

**Figure 3-5.**
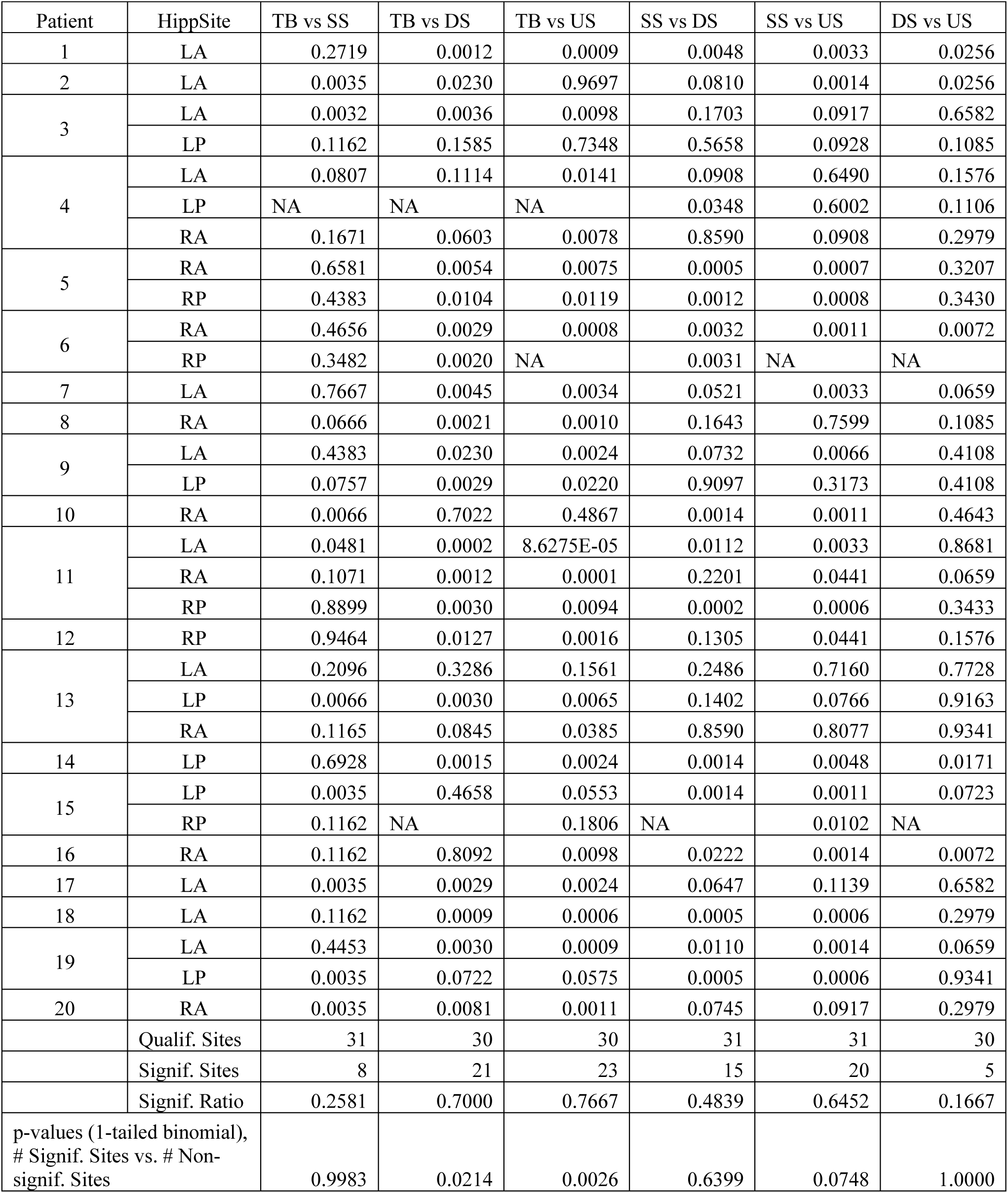
Anatomical distributions of NC sites significantly coupled with HC-SWR in terms of different NC-GE types tend to be similar for some NC-GE type pairs (TB vs. SS, DS vs. US) and different for other pairs. Wilcoxon signed rank tests were performed on the data obtained for Figure 3-3. Each test (whose p-value was tabulated above) required that the corresponding cell in Figure 3-3 for that particular HC site must have significant chi-square test result (thereby counting towards Qualif. Sites); otherwise, the corresponding cell would be filled with NA. # signif.: number of significant tests (in total number of qualified patients).

**Table 3-1.** ROIs for statistical analyses of spatio-temporal differences across NC regions in NC-SWR relationships.

| Combined ROI | Original Freesurfer Label(s) |
| --- | --- |
| Lateral temporal | Banks of superior temporal gyrus, inferior temporal, middle temporal, superior temporal, temporal pole, transverse temporal |
| Cingulate | Caudal anterior cingulate, rostral anterior cingulate, isthmus cingulate, posterior cingulate |
| Orbitofrontal | Frontal pole, lateral orbitofrontal, medial orbitofrontal, pars orbitalis |
| Prefrontal | Caudal middle frontal, pars opercularis, pars triangularis, rostral middle frontal, superior frontal |
| Paracentral | Paracentral, postcentral, precentral |
| Medial temporo-occipital | Entorhinal, fusiform, lingual |
| Medial occipito-parietal | Cuneus, pericalcarine, precuneus |
| Insula | Insula |
| Lateral occipital | Lateral occipital |
| Lateral parietal | Inferior parietal, superior parietal, supramarginal |
| Fronto-central | All labels for the following Combined ROIs: Cingulate, orbitofrontal, prefrontal, paracentral |
| Non-frontal | All labels for the following Combined ROIs: Lateral temporal, medial temporo-occipital, medial occipito-parietal, insula, lateral occipital, lateral parietal |

## References

1. Adler DH et al. (2018) Characterizing the human hippocampus in aging and Alzheimer’s disease using a computational atlas derived from ex vivo MRI and histology. Proc Natl Acad Sci 115:4252–4257.

2. Alekseichuk I, Turi Z, Amador de Lara G, Antal A, Paulus W (2016) Spatial working memory in humans depends on theta and high gamma synchronization in the prefrontal cortex. Curr Biol 26:1513–1521.

3. Andrillon T, Nir Y, Staba RJ, Ferrarelli F, Cirelli C, Tononi G, Fried I (2011) Sleep spindles in humans: insights from intracranial EEG and unit recordings. J Neurosci 31:17821–17834.

4. Axmacher N, Elger CE, Fell J (2008) Ripples in the medial temporal lobe are relevant for human memory consolidation. Brain J Neurol 131:1806–1817.

5. Badre D, Kayser AS, D’Esposito M (2010) Frontal cortex and the discovery of abstract action rules. Neuron 66:315–326.

6. Benjamini Y, Hochberg Y (1995) Controlling the false discovery rate: a practical and powerful approach to multiple testing. J R Stat Soc Ser B Methodol 57:289–300.

7. Bragin A, Engel J, Wilson CL, Fried I, Buzsáki G (1999) High-frequency oscillations in human brain. Hippocampus 9:137–142.

8. Brázdil M, Cimbálník J, Roman R, Shaw DJ, Stead MM, Daniel P, Jurák P, Halámek J (2015) Impact of cognitive stimulation on ripples within human epileptic and non-epileptic hippocampus. BMC Neurosci 16:47.

9. Buzsáki G (2015) Hippocampal sharp wave-ripple: A cognitive biomarker for episodic memory and planning. Hippocampus 25:1073–1188.

10. Buzsaki G, Horvath Z, Urioste R, Hetke J, Wise K (1992) High-frequency network oscillation in the hippocampus. Science 256:1025–1027.

11. Buzsáki G, Logothetis N, Singer W (2013) Scaling brain size, keeping timing: evolutionary preservation of brain rhythms. Neuron 80:751–764.

12. Cairney SA, Guttesen A á V, Marj NE, Staresina BP (2018) Memory consolidation is linked to spindle-mediated information processing during sleep. Curr Biol 28:948–954.e4.

13. Canolty RT, Edwards E, Dalal SS, Soltani M, Nagarajan SS, Kirsch HE, Berger MS, Barbaro NM, Knight RT (2006) High gamma power is phase-locked to theta oscillations in human neocortex. Science 313:1626–1628.

14. Carskadon MA, Dement WC (2010) Monitoring and staging human sleep. In: Principles and practice of sleep medicine, 5th ed. (Kryger MH, Roth T, Dement WC, eds), pp 16–26. St. Louis: Elsevier Saunders.

15. Cash SS, Halgren E, Dehghani N, Rossetti AO, Thesen T, Wang C, Devinsky O, Kuzniecky R, Doyle W, Madsen JR, Bromfield E, Erőss L, Halász P, Karmos G, Csercsa R, Wittner L, Ulbert I (2009) The human K-complex represents an isolated cortical down-state. Science 324:1084–1087.

16. Catenoix H, Magnin M, Mauguière F, Ryvlin P (2011) Evoked potential study of hippocampal efferent projections in the human brain. Clin Neurophysiol 122:2488–2497.

17. Clemens Z, Mölle M, Eross L, Barsi P, Halász P, Born J (2007) Temporal coupling of parahippocampal ripples, sleep spindles and slow oscillations in humans. Brain J Neurol 130:2868–2878.

18. Clemens Z, Mölle M, Eross L, Jakus R, Rásonyi G, Halász P, Born J (2011) Fine-tuned coupling between human parahippocampal ripples and sleep spindles. Eur J Neurosci 33:511–520.

19. Cohen J (1969) Statistical power analysis for the behavioral sciences. New York: Academic Press.

20. Csercsa R et al. (2010) Laminar analysis of slow wave activity in humans. Brain J Neurol 133:2814–2829.

21. Dale AM, Fischl B, Sereno MI (1999) Cortical surface-based analysis. I. Segmentation and surface reconstruction. NeuroImage 9:179–194.

22. de Lavilléon G, Lacroix MM, Rondi-Reig L, Benchenane K (2015) Explicit memory creation during sleep demonstrates a causal role of place cells in navigation. Nat Neurosci 18:493–495.

23. Delorme A, Makeig S (2004) EEGLAB: an open source toolbox for analysis of single-trial EEG dynamics including independent component analysis. J Neurosci Methods 134:9–21.

24. Desikan RS, Ségonne F, Fischl B, Quinn BT, Dickerson BC, Blacker D, Buckner RL, Dale AM, Maguire RP, Hyman BT, Albert MS, Killiany RJ (2006) An automated labeling system for subdividing the human cerebral cortex on MRI scans into gyral based regions of interest. NeuroImage 31:968– 980.

25. Ding S-L, Van Hoesen GW (2015) Organization and detailed parcellation of human hippocampal head and body regions based on a combined analysis of cyto- and chemoarchitecture. J Comp Neurol 523:2233–2253.

26. Duvernoy H (1988) The human hippocampus: an atlas of applied anatomy. Munich, BY: J.F. Bergmann-Verlag.

27. Dykstra AR, Chan AM, Quinn BT, Zepeda R, Keller CJ, Cormier J, Madsen JR, Eskandar EN, Cash SS (2012) Individualized localization and cortical surface-based registration of intracranial electrodes. NeuroImage 59:3563–3570.

28. Ego-Stengel V, Wilson MA (2010) Disruption of ripple-associated hippocampal activity during rest impairs spatial learning in the rat. Hippocampus 20:1–10.

29. Enatsu R, Gonzalez-Martinez J, Bulacio J, Kubota Y, Mosher J, Burgess RC, Najm I, Nair DR (2015) Connections of the limbic network: A corticocortical evoked potentials study. Cortex 62:20–33.

30. English DF, Peyrache A, Stark E, Roux L, Vallentin D, Long MA, Buzsáki G (2014) Excitation and inhibition compete to control spiking during hippocampal ripples: intracellular study in behaving mice. J Neurosci 34:16509–16517.

31. Fischl B, Sereno MI, Dale AM (1999a) Cortical surface-based analysis. II: Inflation, flattening, and a surface-based coordinate system. NeuroImage 9:195–207.

32. Fischl B, Sereno MI, Tootell RB, Dale AM (1999b) High-resolution intersubject averaging and a coordinate system for the cortical surface. Hum Brain Mapp 8:272–284.

33. Fischl B, van der Kouwe A, Destrieux C, Halgren E, Ségonne F, Salat DH, Busa E, Seidman LJ, Goldstein J, Kennedy D, Caviness V, Makris N, Rosen B, Dale AM (2004) Automatically parcellating the human cerebral cortex. Cereb Cortex 14:11–22.

34. Fletcher PC, Henson RNA (2001) Frontal lobes and human memory: insights from functional neuroimaging. Brain 124:849–881.

35. Froemke RC, Dan Y (2002) Spike-timing-dependent synaptic modification induced by natural spike trains. Nature 416:433–438.

36. Genzel L, Kroes MCW, Dresler M, Battaglia FP (2014) Light sleep versus slow wave sleep in memory consolidation: a question of global versus local processes? Trends Neurosci 37:10–19.

37. Gervasoni D, Lin S-C, Ribeiro S, Soares ES, Pantoja J, Nicolelis MAL (2004) Global forebrain dynamics predict rat behavioral states and their transitions. J Neurosci 24:11137–11147.

38. Glosser G, Deutsch GK, Cole LC, Corwin J, Saykin AJ (1998) Differential lateralization of memory discrimination and response bias in temporal lobe epilepsy patients. J Int Neuropsychol Soc 4:502–511.

39. Gonzalez C, Mak-McCully R, Rosen B, Cash SS, Chauvel P, Bastuji H, Rey M, Halgren E (2018) Theta bursts precede, and spindles follow, cortical and thalamic downstates in human NREM sleep. J Neurosci 38:9989–10001.

40. Gonzalez-Martinez J, Bulacio J, Alexopoulos A, Jehi L, Bingaman W, Najm I (2013) Stereoelectroencephalography in the “difficult to localize” refractory focal epilepsy: early experience from a North American epilepsy center. Epilepsia 54:323–330.

41. Halgren E, Kaestner E, Marinkovic K, Cash SS, Wang C, Schomer DL, Madsen JR, Ulbert I (2015) Laminar profile of spontaneous and evoked theta: Rhythmic modulation of cortical processing during word integration. Neuropsychologia 76:108–124.

42. Halgren M, Fabó D, Ulbert I, Madsen JR, Erőss L, Doyle WK, Devinsky O, Schomer D, Cash SS, Halgren E (2018) Superficial slow rhythms integrate cortical processing in humans. Sci Rep 8:2055.

43. Hrybouski S, MacGillivray M, Huang Y, Madan CR, Carter R, Seres P, Malykhin NV (2019) Involvement of hippocampal subfields and anterior-posterior subregions in encoding and retrieval of item, spatial, and associative memories: Longitudinal versus transverse axis. NeuroImage 191:568–586.

44. Ji D, Wilson MA (2007) Coordinated memory replay in the visual cortex and hippocampus during sleep. Nat Neurosci 10:100–107.

45. Jiang X, Gonzalez-Martinez J, Halgren E (submitted-2) Posterior hippocampal spindle-ripples phase-locked with parietal spindles during NREM sleep in humans. unpublished.

46. Jiang X, Shamie I, Doyle W, Friedman D, Dugan P, Devinsky O, Eskandar E, Cash SS, Thesen T, Halgren E (2017) Replay of large-scale spatio-temporal patterns from waking during subsequent NREM sleep in human cortex. Sci Rep 7:17380.

47. Johnson L, Euston D, Tatsuno M, McNaughton B (2010) Stored-trace reactivation in rat prefrontal cortex is correlated with down-to-up state fluctuation density. J Neurosci 30:2650–2661.

48. Latchoumane C-FV, Ngo H-VV, Born J, Shin H-S (2017) Thalamic spindles promote memory formation during sleep through triple phase-locking of cortical, thalamic, and hippocampal rhythms. Neuron 95:424–435.

49. Le Van Quyen M, Bragin A, Staba R, Crépon B, Wilson CL, Engel J (2008) Cell type-specific firing during ripple oscillations in the hippocampal formation of humans. J Neurosci 28:6104–6110.

50. Logothetis NK, Eschenko O, Murayama Y, Augath M, Steudel T, Evrard HC, Besserve M, Oeltermann A (2012) Hippocampal-cortical interaction during periods of subcortical silence. Nature 491:547– 553.

51. Maingret N, Girardeau G, Todorova R, Goutierre M, Zugaro M (2016) Hippocampo-cortical coupling mediates memory consolidation during sleep. Nat Neurosci 19:959–964.

52. Mak-McCully RA, Rolland M, Sargsyan A, Gonzalez C, Magnin M, Chauvel P, Rey M, Bastuji H, Halgren E (2017) Coordination of cortical and thalamic activity during non-REM sleep in humans. Nat Commun 8:15499.

53. Mak-McCully RA, Rosen BQ, Rolland M, Régis J, Bartolomei F, Rey M, Chauvel P, Cash SS, Halgren E (2015) Distribution, amplitude, incidence, co-occurrence, and propagation of human K-complexes in focal transcortical recordings. eNeuro 2:ENEURO.0028-15.2015

54. McCormick DA, Bal T (1997) Sleep and arousal: thalamocortical mechanisms. Annu Rev Neurosci 20:185–215.

55. Mölle M, Yeshenko O, Marshall L, Sara SJ, Born J (2006) Hippocampal sharp wave-ripples linked to slow oscillations in rat slow-wave sleep. J Neurophysiol 96:62–70.

56. Moraes W, Piovezan R, Poyares D, Bittencourt LR, Santos-Silva R, Tufik S (2014) Effects of aging on sleep structure throughout adulthood: a population-based study. Sleep Med 15:401–409.

57. Niethard N, Ngo H-VV, Ehrlich I, Born J (2018) Cortical circuit activity underlying sleep slow oscillations and spindles. Proc Natl Acad Sci 115:E9220–E9229.

58. Niknazar M, Krishnan GP, Bazhenov M, Mednick SC (2015) Coupling of thalamocortical sleep oscillations are important for memory consolidation in humans. PLOS ONE 10:e0144720.

59. Nir Y, Staba RJ, Andrillon T, Vyazovskiy VV, Cirelli C, Fried I, Tononi G (2011) Regional slow waves and spindles in human sleep. Neuron 70:153–169.

60. Oostenveld R, Fries P, Maris E, Schoffelen J-M (2011) FieldTrip: Open source software for advanced analysis of MEG, EEG, and invasive electrophysiological data. Comput Intell Neurosci 2011:156869.

61. Patel J, Schomburg EW, Berényi A, Fujisawa S, Buzsáki G (2013) Local generation and propagation of ripples along the septotemporal axis of the hippocampus. J Neurosci 33:17029–17041.

62. Pavlides C, Winson J (1989) Influences of hippocampal place cell firing in the awake state on the activity of these cells during subsequent sleep episodes. J Neurosci 9:2907–2918.

63. Peyrache A, Khamassi M, Benchenane K, Wiener SI, Battaglia FP (2009) Replay of rule-learning related neural patterns in the prefrontal cortex during sleep. Nat Neurosci 12:919–926.

64. Piantoni G, Halgren E, Cash SS (2017) Spatiotemporal characteristics of sleep spindles depend on cortical location. NeuroImage 146:236–245.

65. Poppenk J, Evensmoen HR, Moscovitch M, Nadel L (2013) Long-axis specialization of the human hippocampus. Trends Cogn Sci 17:230–240.

66. Ramirez-Villegas JF, Logothetis NK, Besserve M (2015) Diversity of sharp-wave–ripple LFP signatures reveals differentiated brain-wide dynamical events. Proc Natl Acad Sci 112:E6379–E6387.

67. Rasch B, Born J (2013) About sleep’s role in memory. Physiol Rev 93:681–766.

68. Rothschild G, Eban E, Frank LM (2016) A cortical-hippocampal-cortical loop of information processing during memory consolidation. Nat Neurosci 20:251–259.

69. Sanchez-Vives MV, McCormick DA (2000) Cellular and network mechanisms of rhythmic recurrent activity in neocortex. Nat Neurosci 3:1027–1034.

70. Sato W, Kochiyama T, Uono S, Matsuda K, Usui K, Inoue Y, Toichi M (2014) Rapid, high-frequency, and theta-coupled gamma oscillations in the inferior occipital gyrus during face processing. Cortex 60:52–68.

71. Schreiner T, Doeller CF, Jensen O, Rasch B, Staudigl T (2018) Theta phase-coordinated memory reactivation reoccurs in a slow-oscillatory rhythm during NREM sleep. Cell Rep 25:296–301.

72. Schreiner T, Rasch B (2015) Boosting vocabulary learning by verbal cueing during sleep. Cereb Cortex 25:4169–4179.

73. Seibt J, Richard CJ, Sigl-Glöckner J, Takahashi N, Kaplan DI, Doron G, Limoges D de, Bocklisch C, Larkum ME (2017) Cortical dendritic activity correlates with spindle-rich oscillations during sleep in rodents. Nat Commun 8:684.

74. Siapas AG, Wilson MA (1998) Coordinated interactions between hippocampal ripples and cortical spindles during slow-wave sleep. Neuron 21:1123–1128.

75. Silber M, Ancoli-Israel S, Bonnet M, Chokroverty S, Grigg-Damberger M, Hirshkowitz M, Kapen S, Keenan S, Kryger M, Penzel T, Pressman M, Iber C (2007) The visual scoring of sleep in adults. J Clin Sleep Med 3:121–131.

76. Skaggs WE, McNaughton BL, Permenter M, Archibeque M, Vogt J, Amaral DG, Barnes CA (2007) EEG sharp waves and sparse ensemble unit activity in the macaque hippocampus. J Neurophysiol 98:898–910.

77. Squire LR, Clark RE, Knowlton BJ (2001) Retrograde amnesia. Hippocampus 11:50–55.

78. Staba RJ, Wilson CL, Bragin A, Jhung D, Fried I, Engel J (2004) High-frequency oscillations recorded in human medial temporal lobe during sleep. Ann Neurol 56:108–115.

79. Staresina BP, Bergmann TO, Bonnefond M, van der Meij R, Jensen O, Deuker L, Elger CE, Axmacher N, Fell J (2015) Hierarchical nesting of slow oscillations, spindles and ripples in the human hippocampus during sleep. Nat Neurosci 18:1679–1686.

80. Steriade M (2001) Impact of network activities on neuronal properties in corticothalamic systems. J Neurophysiol 86:1–39.

81. Strange BA, Witter MP, Lein ES, Moser EI (2014) Functional organization of the hippocampal longitudinal axis. Nat Rev Neurosci 15:655–669.

82. Suh J, Foster DJ, Davoudi H, Wilson MA, Tonegawa S (2013) Impaired hippocampal ripple-associated replay in a mouse model of schizophrenia. Neuron 80:484–493.

83. Suzuki WA, Amaral DG (2004) Functional neuroanatomy of the medial temporal lobe memory system. Cortex 40:220–222.

84. Van Hoesen GW (1995) Anatomy of the medial temporal lobe. Magn Reson Imaging 13:1047–1055.

85. Wilson MA, McNaughton BL (1994) Reactivation of hippocampal ensemble memories during sleep. Science 265:676–679.

86. Zhang H, Fell J, Axmacher N (2018) Electrophysiological mechanisms of human memory consolidation. Nat Commun 9:4103.

